# Beyond Direct Binding: Serotonin Rewires Model Brain Plasma Membrane-Mediated Communication Pathways in Y4R

**DOI:** 10.64898/2026.10.31.763150

**Authors:** Somdatta Paul, Pooja Sharma, Samapika Sahu, Ananya Debnath

## Abstract

Serotonin, an endogenous neurotransmitter, modulates pancreatic polypeptide (PP) recognition by the neuropeptide Y4 receptor (Y4R), a mechanism relevant to altered serotonergic signaling associated with metabolic disorder and anxiety. Although recent experiment demonstrates serotonin modulated PP dissociation kinetics of Y4R through membrane allostery, the molecular mechanism of serotonin-induced lipidomic reorganization of the brain plasma membrane (BPM) relevant for Y4R activation is unknown. To this end, a total of 825.9 ns long all-atom molecular dynamics simulations at 310 K elucidate the concentration-dependent coupling between serotonin partitioning, selective lipid reorganization, and PP-binding and Y4R activation relevant intra-receptor communication in a heterogeneous, asymmetric model BPM. In the absence of serotonin, Y4R preferentially associates with the anionic lipids. Serotonin preferentially localizes at the lipid–water interface over the PP-binding and activation-associated regions of Y4R. At 6 mM, serotonin preferentially contacts phosphocholine species depleted from Y4R, whereas at 50 mM, phosphatidylinositol 4,5-bisphosphate dominates both micro-environments. These changes align with the selective remodeling of lipid networks, enhanced lipid-chain ordering, and modest bilayer compaction. Weighted implementation of suboptimal paths analysis reveals concentration-dependent reorganization of communication pathways between experimentally implicated PP-binding and activation-associated Y4R residues. Collectively, these findings connects serotonin dependent selective lipid redistribution to altered intramolecular communication between Y4R residues associated with PP binding and G-protein coupling, and provide a framework for future experiments investigating membrane-mediated neuromodulation and neuronal signaling. These findings provide a basis for exploring lipid-targeted pharmacology as a complementary strategy to modulate GPCR allostery by altering membrane composition rather than directly targeting the receptor.

## Introduction

For decades, rational drug design has been guided predominantly by the classical “lock-and-key” paradigm, emphasizing ligand recognition at orthosteric or allosteric sites of target proteins^1^. However, receptor function depends not only on protein structure and ligand binding but also on the surrounding plasma membrane (PM), which regulates receptor organization, conformational dynamics, and signaling^2^. This membrane-centric perspective is particularly relevant to neuropharmacology, where peripheral hormonal signals interact with neuronal receptors to coordinate metabolic and behavioral responses.

The pancreas–brain axis exemplifies this interplay between peripheral metabolism and central signaling. Pancreatic polypeptide (PP), released by the pancreas, activates neuropeptide Y receptor Y4 (Y4R), a class A G protein-coupled receptor (GPCR)^3^ that transduces extracellular signals through intracellular G proteins. Y4R–PP signaling contributes to appetite regulation and energy homeostasis and has also been implicated in anxiety-related behaviors^4^. Understanding how the neuronal membrane environment influences this receptor–ligand interaction may therefore provide insights into the integration of metabolic and behavioral signaling.

Serotonin (5-hydroxytryptamine, 5-HT) is an endogenous amphiphilic neurotransmitter that primarily signals through its cognate serotonin receptors, most of which are GPCRs. Altered serotonergic signaling is associated with psychiatric disorders, including depression and anxiety, while also contributing to appetite regulation and energy homeostasis^5–8^. Although 5-HT itself is not a drug, serotonergic medications and psychedelics can modify serotonergic signaling and may also influence membrane organization^9^. Importantly, 5-HT may also modulate non-cognate GPCR signaling indirectly through changes in membrane lipid organization and receptor–membrane interactions. Recent fluorescence spectroscopy, atomic force microscopy (AFM), and solid-state nuclear magnetic resonance (NMR) studies indicate that increasing 5-HT concentrations are associated with a decrease in membrane order and an increase in the dissociation constant of the Y4R–PP complex for a particular lipid composition^10^. These findings suggest that serotonin-induced membrane remodeling may modulate Y4R–PP interactions, although the underlying molecular mechanisms remain unresolved. More broadly, experimental and computational studies have established that small molecules and drugs can alter membrane properties, highlighting the need to understand how such changes influence receptor signaling^11^.

Neuronal plasma membranes contain hundreds of lipid species whose composition and spatial distribution regulate membrane organization and cellular function^12^. Cholesterol (CHOL) and sphingomyelin (SM) contribute to dynamic nanoscale membrane domains, commonly termed lipid rafts, that organize signaling components and influence receptor behavior^13,14^. Polyunsaturated fatty acid (PUFA)-rich glycerophospholipids modulate membrane fluidity and flexibility, whereas phosphoinositides (PIPs) regulate membrane surface charge, trafficking, and protein interactions^15^. Consequently, variations in lipid composition, membrane thickness, stiffness, elasticity, and curvature can alter GPCR stability, conformational dynamics, and activity^16^. Stress-associated changes in lipid composition, including lipid peroxidation and altered PUFA ratios, may further disrupt membrane organization and protein–lipid interactions, with potential implications for psychiatric and neurodegenerative conditions^17,18^. Elucidating how lipid heterogeneity shapes neuronal membrane biophysics is therefore important for understanding receptor regulation under physiological and stress-related conditions.

GPCRs and their surrounding lipids exhibit reciprocal coupling: receptors can redistribute membrane lipids and alter lateral packing, surface tension, and domain organization, while the local lipid environment influences receptor conformation and signaling independently of direct ligand binding^19^. Despite this growing evidence, the molecular basis of membrane-mediated modulation of non-cognate GPCR signaling remains poorly understood. Here, Y4R provides a relevant model for investigating whether changes in neuronal membrane lipid organization alter PP recognition and receptor-associated interactions. By examining the interplay between membrane lipid composition, 5-HT concentration, and Y4R–PP association, this work extends the conventional protein-centric view of receptor pharmacology and explores the neuronal membrane as an active regulator of pancreas–brain signaling, with potential implications for membrane-targeted therapeutic strategies.

Thus, the objective of the current work is to find out: a) how does serotonin reshape the heterogeneous brain membrane lipid landscape to modulate non-cognate Y4R–PP interactions in the absence of direct serotonin–receptor contact? b) How does serotonin-induced lipid reorganization rewire intrareceptor communication pathways in Y4R while preserving its overall transmembrane architecture? To this end, a total of 825.9 ns long AA MD simulations are performed for a transmembrane Y4R-PP complex embedded in a heterogeneous model brain plasma membrane^20^ at three different 5-HT concentrations at 310 K using the CHARMM36m force field. Across five independent molecular dynamics replicas per condition, our simulations reveal a concentration-dependent, membrane-mediated mechanism for serotonergic modulation of the Y4R–PP complex. The 6 mM 5-HT condition is selected to relate our simulations with previous experimental evidence of serotonin-mediated modulation of PP–Y4R binding kinetics^10^, while the 50 mM condition is included to probe how a higher serotonin concentration influences membrane organization and receptor-associated communication. Consistent with the experimental observation that 5-HT can modulate Y4R–PP interactions without direct receptor contact^10^, our simulations show that 5-HT preferentially localizes at the lipid–water interface and promotes selective lipid redistribution. Y4R retains a preferential association with the anionic lipids, particularly PIP2 followed by PS, while increasing 5-HT concentration drives biphasic PIP2 sorting and selectively remodels the PIP2–PS interaction network, whereas the CER–SM-associated network remains comparatively conserved. Unlike the generalized membrane disordering reported for single-component membranes, the heterogeneous model brain PM exhibits modest lipid-chain ordering and membrane compaction with increasing 5-HT concentration. Most notably, weighted implementation of suboptimal paths (WISP) analysis reveals concentration-dependent rewiring of intra-receptor communication between the experimentally determined PP-binding residue TYR101^21^ and activation residue ASP289 and ARG139 of Y4R^21,22^. The TYR101-ASP289 communication path is dominated by TRP107 at 0 mM from our calculations. The same residue is known for PP recognition and Y4R activation from the previous experimental observation^21^. The path is replaced by alternative communication routes at 6 mM, while at 50 mM the principal route is broadly preserved but exhibits reduced node degeneracy and greater contributions from alternative residues. Similarly, the communication pathway of the TYR101-ARG139 of Y4R, residing on two opposing leaflets of the membrane, is modulated via altered node degeneracy of involved residues due to 5-HT. These changes coincide with altered fluctuations and tilt angles of implicated receptor elements. Together, these findings connect serotonin-dependent membrane lipid reorganization and preferential Y4R–PIP2/PS association with changes in intra-receptor communication of Y4R that involves PP binding and signaling residues. The results provide an atomistic framework consistent with experimentally observed membrane-mediated modulation of Y4R–PP interactions. Overall, the work establishes a membrane-centric framework for non-cognate GPCR regulation, opening new avenues for lipid-targeted drug discovery and precision therapeutics for mental disorders^23^.

## Results and Discussion

### 5-HT Prefers to Reside near the Lipid Head Groups over the Signaling Residues of Y4R-PP for All Concentrations

The main core of the Y4R protein, as shown in Figure 1A, comprises a bundle of seven transmembrane (TM) helices (TM1–TM7) that span the hydrophobic lipid bilayer, physically connecting the extracellular environment to the cytosolic side. The receptor utilizes multiple flexible linkers, specifically the extracellular and cytosolic loops, to interconnect these TM domains. Additionally, the protein has a helix-8 (H8), which does not cross the membrane; it resides entirely within the aqueous cytosolic region, functioning as a primary interaction site for downstream cellular signaling^22^. Additional structural features of Y4R include short *β*- sheet segments within the extracellular domain, an N-terminus, and a C-terminus, which reside outside the trans-membrane region and are solvated in water. The cognate ligand, PP, is comprised of a central *α*-helix (A1) and unstructured N and C termini that interact predominantly with the extracellular-facing vestibule of the trans-membrane pocket. It is known that the top of the TM2, TM6, and TM7 are the preferred binding pocket for the PP ligand^21^.

**Figure 1.**
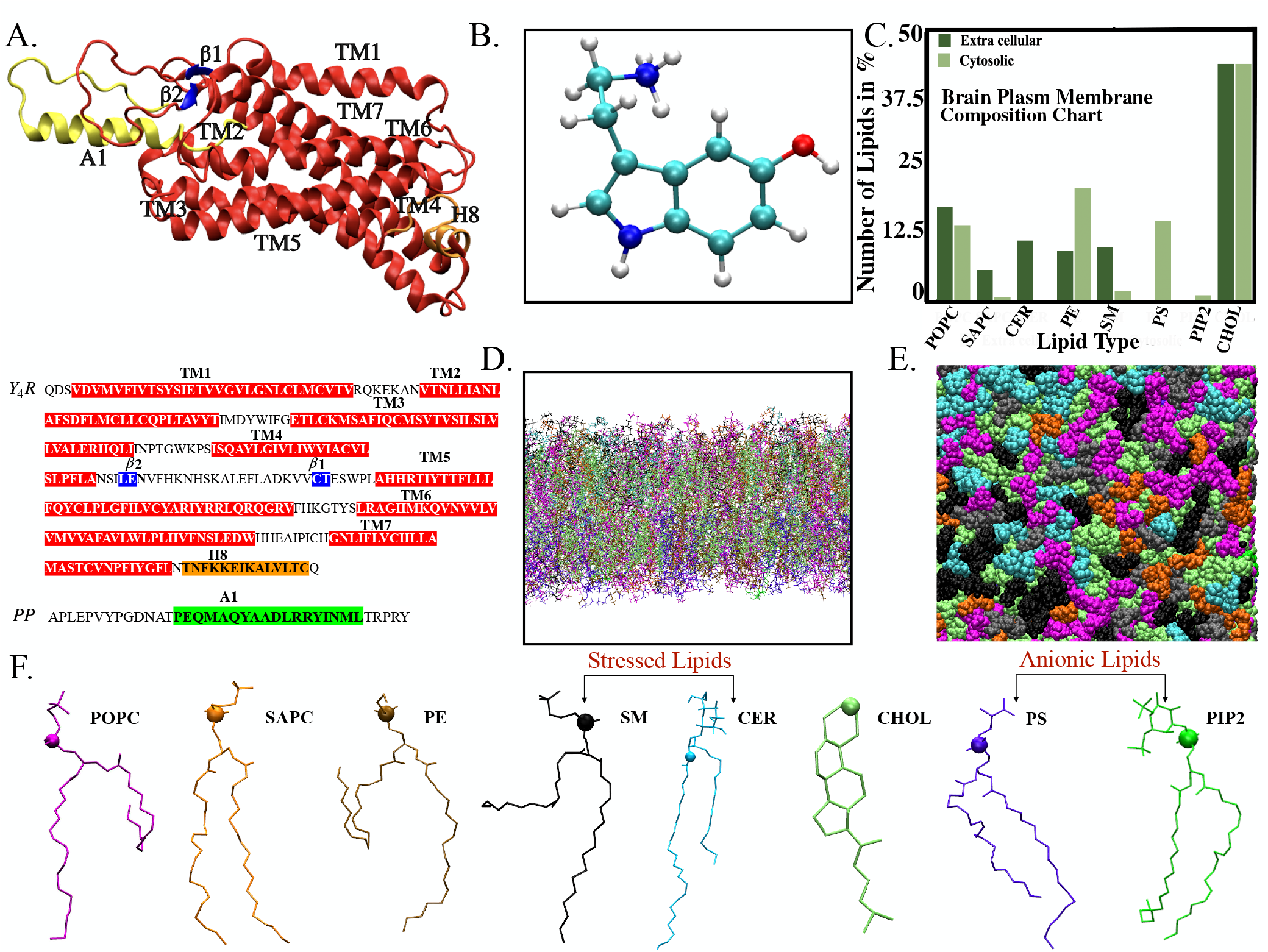
(A) Snapshot of the Y4R-PP showing PP ligand (yellow), seven trans-membrane helices (TM 1–7), and helix 8 (H8) of Y4R (red). Two small *β*-sheets are shown in blue. The amino acid sequences of Y4R and PP used in the simulation start from Gln35 and end at Gln341. (B) Three-dimensional ball-and- stick representation of the geometry-optimized neurotransmitter protonated 5-HT. (C) Quantitative lipid composition profile detailing the asymmetric distribution of distinct lipid species (POPC, SAPC, CER, PE, SM, PS, PIP, and CHOL) across the extracellular (dark green) and cytosolic (light green) leaflets. (D) Side-view of the AA brain plasma membrane bilayer. (E) Top-view of the equilibrated brain plasma membrane illustrating the lateral heterogeneity and complex packing of the diverse lipid components. (F) Ball-and- stick representation of each lipid species. Color code: POPC: magenta, SAPC: orange, PE: ochre, SM: black, CHOL: lime, PS: violet, PIP2: green.

The initial configuration of 5-HT is shown in Figure 1 B. The membrane bilayer is designed using the simplified, asymmetric, model Brain Plasma Membrane (BPM), developed by Ingólfsson et al.^20^, derived from quantitative mammalian lipidomic data which captures the essential biophysical properties and lipid asymmetry conserved in human neuronal membranes. Hereafter, this setup is referred to as the model brain plasma membrane (BPM). The asymmetric membrane comprises a heterogeneous mixture of eight lipid species. The percentage of lipids in both leaflets is mentioned in Figure 1 C. CHOL is the most abundant lipid in both leaflets, followed by POPC, PE, PS, CER, and SM. A small percentage of PIP2 lipid is present only in the leaflet facing the cytosolic side. PS is present only in the same leaflet. CER is only present in the other leaflet. The lipid heterogeneity is also noticeable from the Figure 1 D-E. The chemical structures of the constituent lipid species are illustrated in 1F, highlighting their structural diversity, which underlies their distinct biophysical roles in modulating membrane organization, fluidity, and receptor interactions.

The final configurations of the production runs of the bilayers simulated at three 5-HT concentrations are shown in Figure 2. For all concentrations, 5-HT is located at the lipid-water interface (Figure 3). The peaks of the density distributions of 5-HT and most of the lipid heads share similar locations (Figure 3). 5-HT, preferentially localizes near the lipid headgroups, more specifically to the PC and anionic lipids (PS, PIP2), rather than penetrating deeply into the hydrophobic core of the membrane. Such positioning is consistent with the amphiphilic character of 5-HT, which enables it to interact with both lipid headgroups and the aqueous environment in simulation. The protein density spans across the hydrophobic region of the bilayer, confirming that it is transmembrane in nature and is stably inserted within the lipid environment. The ligand density strongly overlaps with that of the extracellular part of the protein that resides in the extra-cellular side of the bilayer. This suggests favorable interactions of the ligand with the protein and the water molecules.

**Figure 2.**
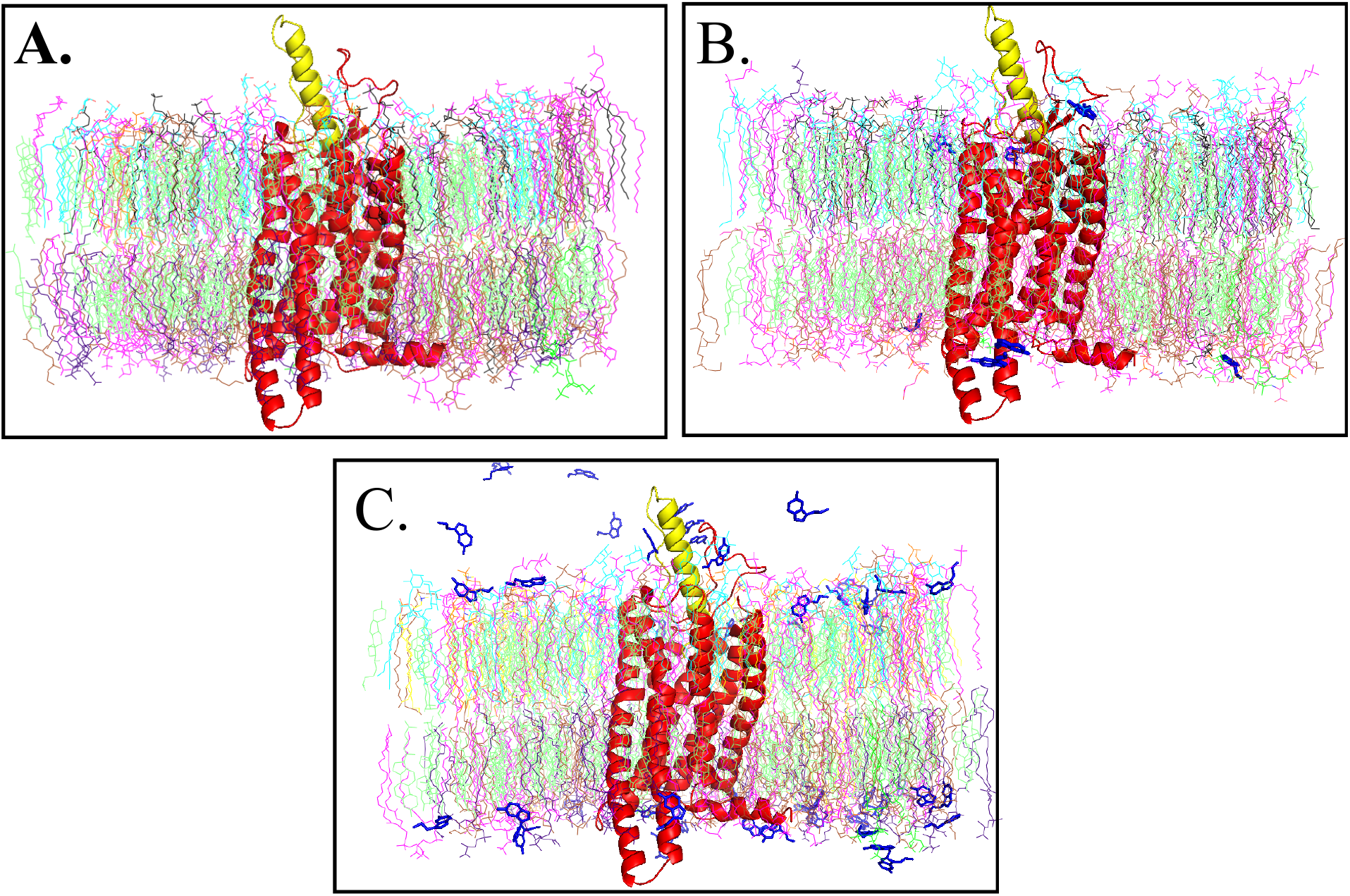
Final snapshots of the Y4R-PP complex embedded in the brain plasma membrane across A) 0, B) 6, and C) 50 mM concentrations. Color code and representation: Y4R: red, new-cartoon representation; bound PP ligand: yellow, new-cartoon; 5-HT: iceblue, licorice, TIP4P/2005 water molecules and neutralizing ions are not shown for visual clarity. The snapshots illustrate the concentration-dependent partitioning and preferential accumulation of 5-HT from S1-S3 at the lipid–water interfacial regions.

**Figure 3.**
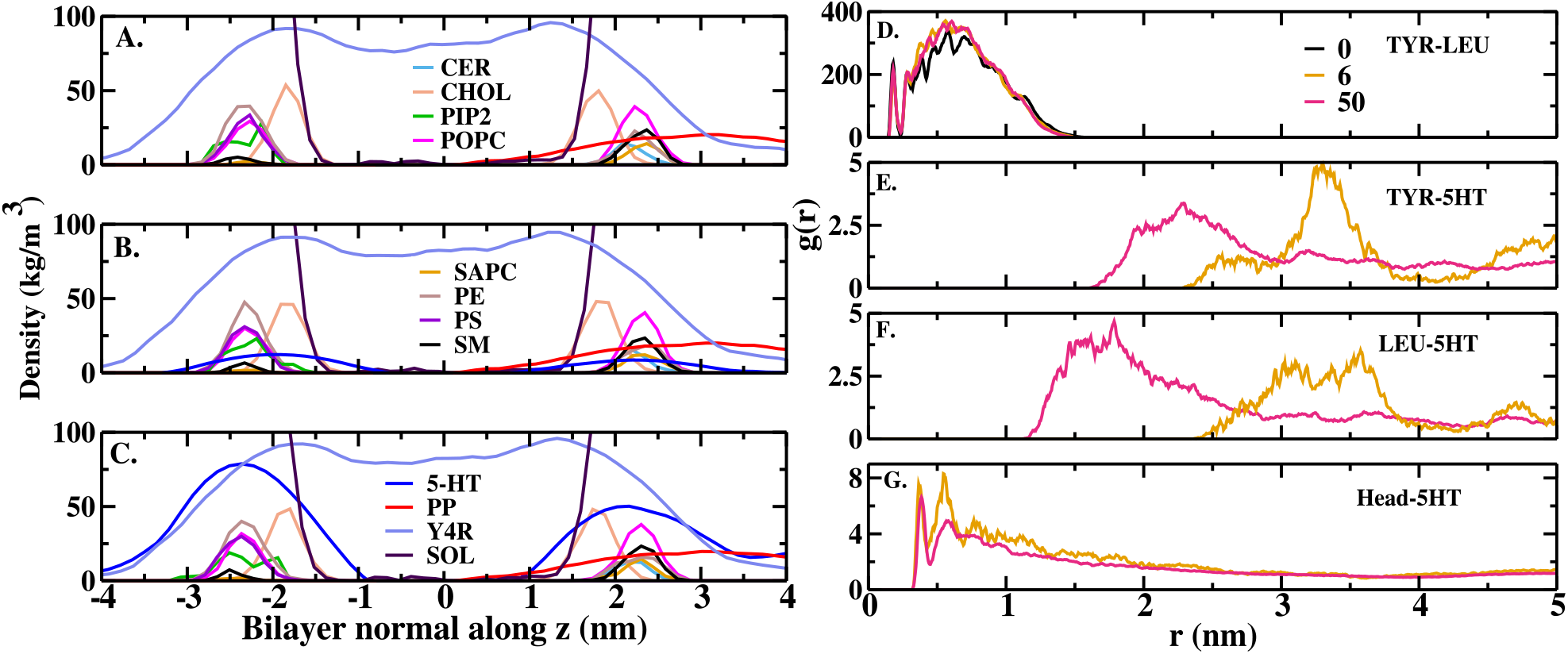
Density profiles along the bilayer normal for systems A) S1, 0 mM, B) S2, 6 mM, and C) S3, 50 mM. The profiles illustrate the spatial distribution and membrane partitioning of the protein (Y4R), ligand (PP), solvent (SOL), 5-HT, and diverse lipid species across leaflets. For visualization, the density values of SERT and PIP2 are scaled by factors of 20 and 10, respectively. Three dimensional (3D) radial distribution function between D) TYR101-LEU31 for three different concentrations and E) TYR101-5-HT F) LEU31-5-HT G) lipid head-5-HT for 6 mM and 50 mM serotonin. This shows 5-HT resides near the lipid head group compared to the receptor or the ligand.

To quantify the spatial preference of 5-HT to Y4R-PP complex over 5-HT, radial distribution functions (RDF), *g*(*r*), are calculated across all the concentrations of serotonin. TYR-101 of helix II of Y4R and LEU-31 of the PP ligand helix A are known to form key contact hubs for signaling^21,24^, without known direct interaction or binding affinity with 5-HT^10^. Upon comparing the first peak of *g*(*r*) of TYR-LEU to TYR-5HT and LEU-5-HT (Figure 3D-F), it is found that 5-HT does not have a direct association with either TYR-101 or LEU-31 of Y4R and PP. Conversely, the RDF between the lipid headgroups and 5-HT (Figure 3G) shows a sharp, highly pronounced first-shell coordination peak. Similarly, the RDF of 5-HT is calculated with respect to the ASP289 from the TM6 of Y4R and ARG139 from the E/DRY motif of Y4R, which are well known for receptor activation through PP recognition or through G-protein binding. The first peak of the RDF (in the Figure S2 (A-C) of the SI) confirms a clear preference of 5-HT for the lipid headgroup environment compared to ARG139 or ASP289 of Y4R. Overall, these RDF analyses confirm that across all concentrations, 5-HT preferentially partitions into the lipid headgroup interface rather than maintaining close spatial proximity to the receptor residue, TYR-101, PP residue, LEU-31, ARG139 or ASP289.

### Zwitterionic Microdomain Partitioning and Biphasic PIP2 Sorting for Y4R and 5-HT Preferences

To elucidate the preferential binding of 5-HT and Y4R to specific lipids, lipid Depletion–Enrichment (D-E) index around the protein is calculated. This identifies which lipid species is preferentially recruited to the local environment of the protein or serotonin relative to the respective species bulk membrane abundance. Across all three 5-HT concentrations, PIP2 exhibits the highest D-E index indicating strong enrichment around the contact shell of the receptor Y4R relative to its overall membrane composition (Figure 4A). The preference decreases in descending order by PS, PE, SM, CER, CHOL, and POPC, with SAPC showing the lowest enrichment or highest depletion among the lipid species examined. Notably, CER and CHOL show D-E index values close to 1 suggesting no preferential interactions. POPC and SAPC have D-E index below 1, indicating depletion by the protein. This enrichment ranking remains largely consistent across the three 5-HT concentrations (0, 6, and 50 mM), suggesting that most of the lipid preference of Y4R is not directly altered by the presence of 5-HT. A modest concentration-dependent increase in D-E index from depletion to no-interaction is observed for SAPC at 50 mM. The protein-lipid contact frequency, which is shown in Figure 4B, closely mirrors the trend of D-E index by showing the highest contact with PIP2 followed by the similar trend. The agreement between the two metrics confirms that PIP2’s enrichment reflects a genuine preferential interaction with the protein rather than simply its relative abundance in the membrane. As with the D-E index, contact frequencies for each lipid species remain largely stable across the three concentrations, with only minor variations, suggesting that the protein’s local lipid environment is not strongly perturbed by increasing 5-HT concentration.

**Figure 4.**
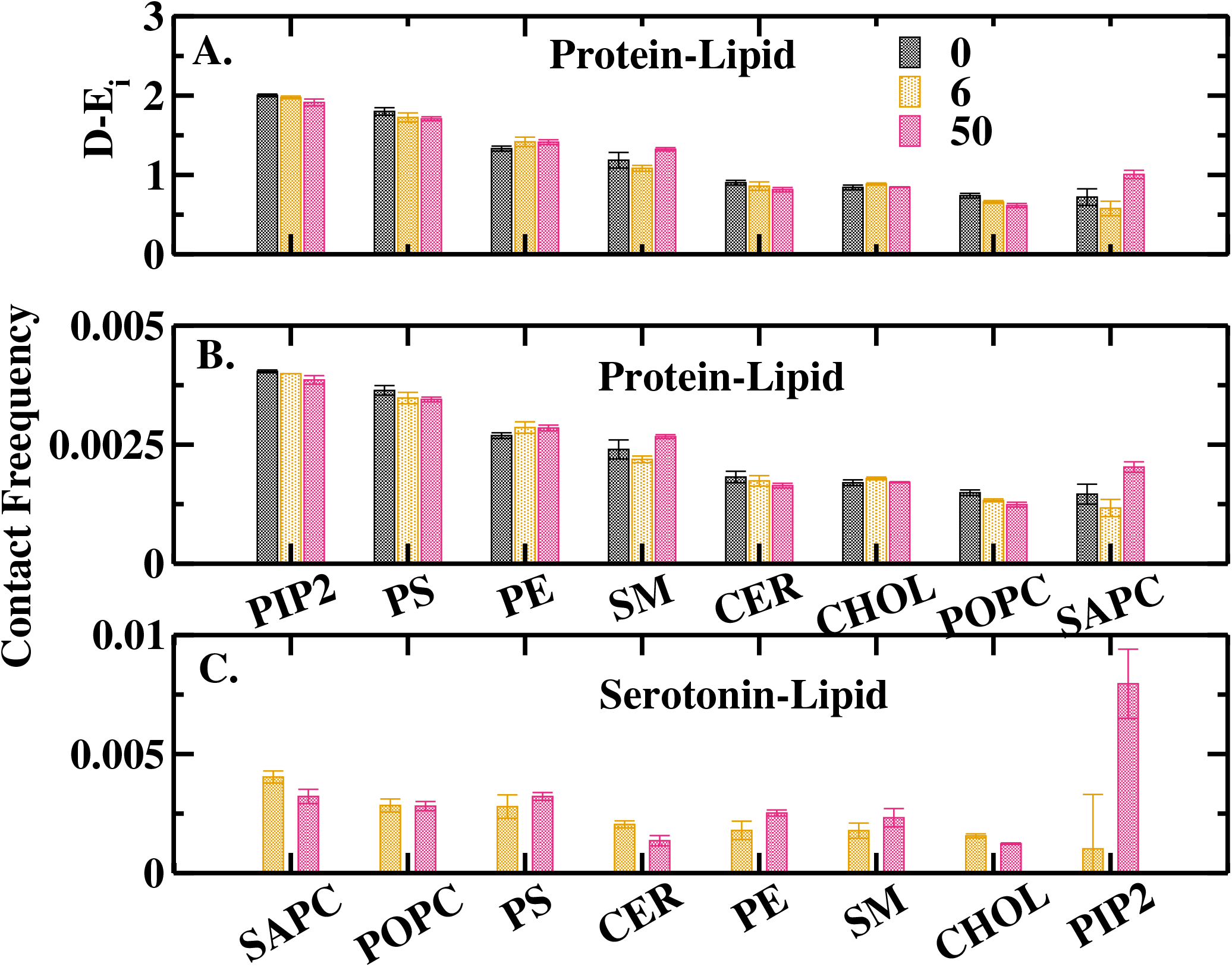
Lipid depletion-enrichment index (D-E_*i*_), protein–lipid and 5-HT–lipid contact frequency across varying 5-HT concentrations.(A) D-E_*i*_ calculated for individual lipid species within the contact shell of the Y4R receptor relative to the bulk bilayer composition at 0 mM, 6 mM, and 50 mM 5-HT concentrations. Values *>* 1 indicate local enrichment, while values *<* 1 denote depletion. (B) Protein–lipid contact frequency across lipid types, mirroring the relative enrichment trends observed in panel A. (C) 5-HT–lipid contact frequency, highlighting a concentration-dependent shift toward PIP_2_ at high 5-HT concentration.

Remarkably, the preferential contacts of serotonin to zwitterioninc PC species are particularly opposite to that of the protein for both concentrations of serotonin. Zwitterionic PC lipids that are depleted from Y4R have the highest contact with 5-HT at both 6 and 50 mM (Figure 4C). This persistent segregation suggests that 5-HT partitions into distinct zwitterionic microdomains, orthogonal to the protein’s native lipid environment. Similarly, PIP2 which prefers to be closer to Y4R, has the least contact with 5-HT at 6 mM. However, the behavior changes with elevated concentration of 5-HT. At 50 mM, a striking shift occurs. 5-HT-PIP2 contacts surge dramatically, making PIP2 the primary lipid species interacting with 5-HT despite its low overall abundance. Thus, the comparison between the protein–lipid and 5-HT–lipid contact frequencies (Figure 4B and C) at 50 mM, reveals that, at 0 and 6 mM, PIP2 is predominantly localized within the immediate boundary of the receptor shell, maintaining the highest protein–lipid and the least serotonin-lipid contact frequency. As 5-HT concentration increases to 50 mM, protein–PIP2 contacts exhibit a modest decline compared to the large upshift of contacts near 5-HT. Since the local accumulation of PIP2 surrounding 5-HT far outpaces its depletion from the receptor interface, PIP2 emerges as the dominant, mutually preferred species within both the receptor and 5-HT microenvironments at 50 mM. As the concentration of 5-HT increases from 6 to 50 mM, the other anionic lipid PS, reduces its contact with the Y4R while simultaneously increasing its contact with 5-HT by a comparable margin.

### 5-HT induces Selective Remodeling of the Anionic-Lipid Associated Network, Keeping the Stressed-Lipid associated Network Comparatively Conserved

To elucidate how the 5-HT concentration lipid–lipid associations, enrichment-based lipid interaction networks are constructed for all concentrations (5). At 0 mM 5-HT (Figure 5A), the membrane exhibits a heterogeneous lipid network, with distinct associations among the major phospholipids (PC), sphingolipid (SM, CER), and sterol components (CHOL). PIP2 is primarily associated with PS and PE, whereas CER and SM form a connected region with CHOL, POPC, and SAPC. This organization represents the baseline of lipid–lipid connectivity in the absence of 5-HT. At 6 mM 5-HT (Figure 5 B), the POPC and SAPC network connectivity disappears, and the PS-CHOL network gets connected. This reorganization is consistent with the contact analysis showing that 5-HT preferentially reside near SAPC, POPC, and PS at 6 mM as shown in Figure 4. Thus, the moderate 5-HT concentration appears to primarily reorganize the connectivity among membrane lipids, whereas the receptor’s overall lipid preference remains unaltered. At 50 mM 5-HT (Figure 5 C), the broken SAPC-POPC network is reformed with a more pronounced organization of the anionic lipid network (PIP2-PS). PIP2 remains strongly associated with PS, while its association to PE is lost, resulting in a more PS-centered network. This organization coincides with the marked increase in 5-HT–PIP2 contacts observed at this concentration (Figure 4C), suggesting that the enhanced interaction of 5-HT with PIP2 is accompanied by remodeling of the local lipid–lipid association network. Importantly, this does not indicate complete displacement of PIP2 from the receptor, as protein–PIP2 contacts remain relatively high (as in Figure 4B), but rather suggests that increasing 5-HT concentration alters the organization and connectivity of PIP2 within the surrounding lipid or membrane micro-environment. For all concentrations of 5-HT, the CHOL–CER-SM-POPC-associated network remains conserved. CER maintains associations with CHOL, POPC, and SM, while SM remains connected with POPC and SAPC. Although their direct contacts with 5-HT vary with concentration, the persistence of these lipid–lipid associations suggests that the SM-associated network is relatively resistant to the concentration-dependent remodeling observed in the PIP2/PS region. Given the established role of CER as a bioactive sphingolipid involved in membrane organization and cellular stress-associated signaling^25^, this conserved CER–SM–CHOL connectivity may represent a structurally persistent component of the membrane rather than a direct signature of 5-HT-induced stress. In addition, increasing 5-HT concentration is associated with selective remodeling of lipid–lipid connectivity, with the most pronounced changes within the PIP2-PS-associated network, while the CER-SM-associated network remains comparatively conserved. Together with the contact-frequency analysis, the lipid network remodeling suggests that 5-HT modulates the receptor-associated membrane environment through selective redistribution of lipids and enhanced engagement of anionic PIP2-containing lipid networks.

**Figure 5.**
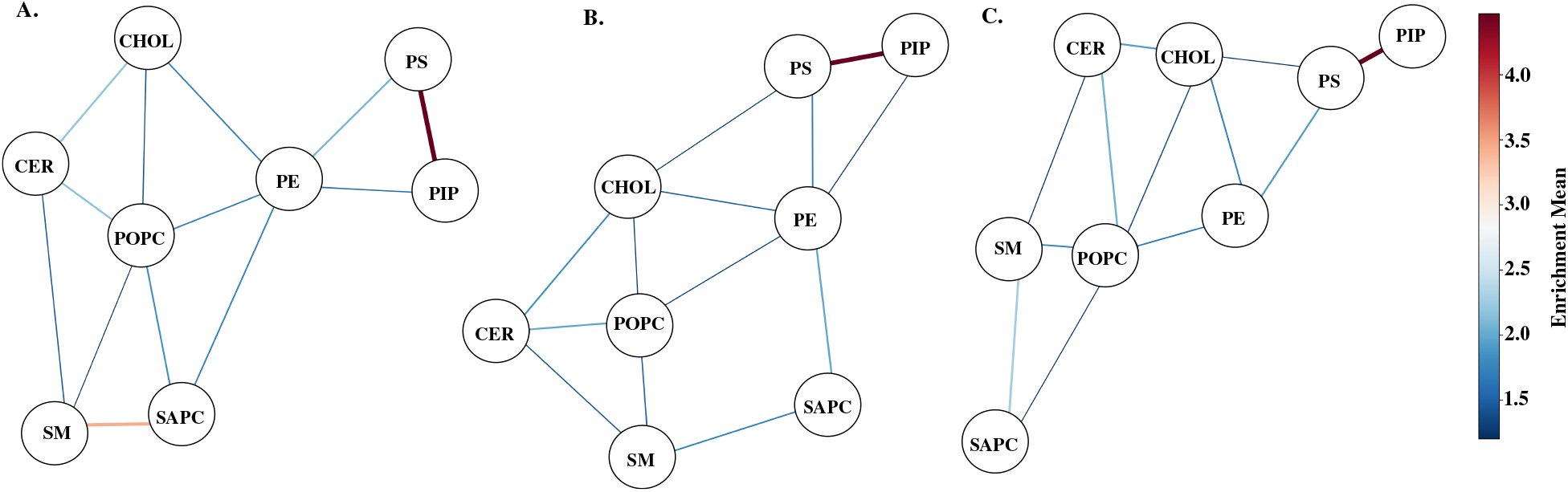
Lipid–lipid enrichment networks at (A) 0 mM, (B) 6 mM, and (C) 50 mM 5-HT. Nodes represent lipid species, and edges represent enriched pairwise lipid associations. Edge width and color indicate the magnitude of the mean enrichment, with the color scale shown at right. The lipid network reveals concentration-dependent reorganization of lipid redistribution in membrane.

### 5-HT-induced Lipid Redistribution by Enhancing Lipid Chain Order

To quantify the effect of 5-HT on membrane lipid packing, the deuterium order parameter (*S*_*CH*_ ) is calculated for each lipid species for three concentrations of 5-HT (Figure 6A,B,C). In Figure 6A it is shown that the individual lipid species exhibit distinct levels of chain ordering, reflecting their individual packing characteristics within the heterogeneous brain plasma membrane. CER and SM exhibit the highest *S*_*CH*_ values, followed by POPC, SAPC, PIP2, PS, PE, and CHOL. Across increasing 5-HT concentrations, most of the individual lipid species show minor increment in *S*_*CH*_, with no systematic decrease in chain ordering. POPC increases slightly from approximately from 0 mM to 50 mM, while CER remains similar across all concentrations. SM shows a small increase at 6 mM and remains nearly unchanged at 50 mM. Similarly, SAPC exhibits a minor increase, whereas PIP2, PS, PE, and CHOL remain comparatively stable. The overall *S*_*CH*_ for all lipids increases with increasing concentrations of 5-HT(Figure 6B). Although the magnitude of this change is not very significant, the progressive increase indicates that the membrane-averaged lipid-chain ordering is not reduced upon 5-HT incorporation. The leaflet wise analysis further demonstrates that (Figure 6C) the two leaflets exhibit somewhat different responses. In the extra-cellular leaflet, *S*_*CH*_ increases progressively from S1 to S2 to S3. In contrast, the *S*_*CH*_ of the cytosolic leaflet remains nearly unchanged from S1 to S2 to S3. Thus, the membrane does not exhibit a generalized disordering response upon increasing 5-HT concentration in contrary to that observed for the single component PC lipids in the earlier experiments. Previous solid-state ^2^H NMR and atomic force microscopy (AFM) studies of a model POPC membrane report that incorporation of 5-HT decreases the acyl-chain order parameter and membrane stiffness^10^. This contrasting response highlights the importance of lipid composition in determining the membrane response to 5-HT. A single-component POPC bilayer represents one specific lipid environment and therefore cannot necessarily predict the ordering behavior of the heterogeneous brain plasma membrane. In contrast, the present model brain plasma membrane contains eight chemically distinct lipid species with different chemical structure and intrinsic packing properties, together with an embedded transmembrane receptor. Consequently, the membrane-averaged *S*_*CH*_ represents the collective response of a structurally heterogeneous lipid ensemble rather than the response of an individual lipid species.

**Figure 6.**
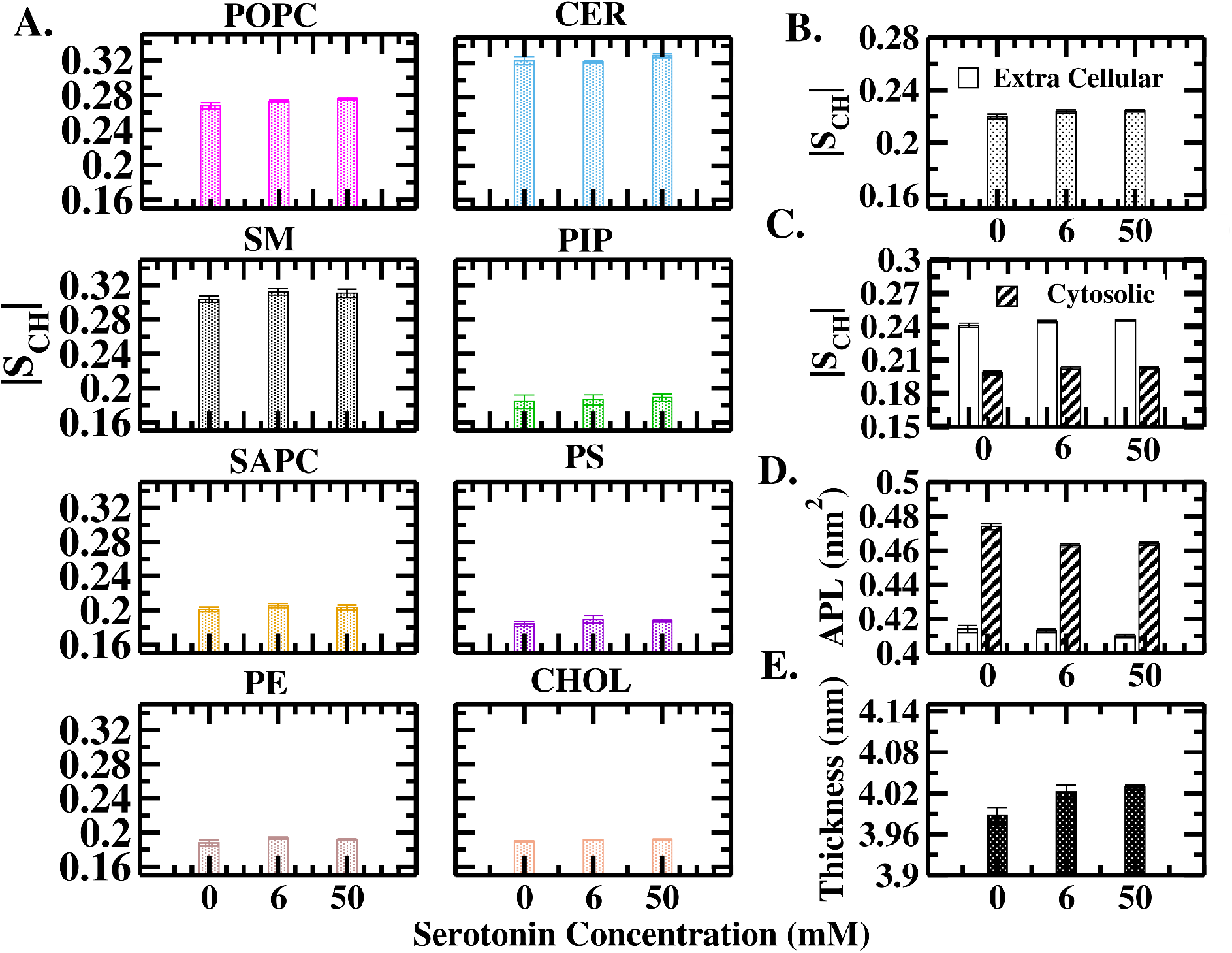
(A) Lipid acyl chain order parameters calculated for individual lipid species (POPC, SM, SAPC, PE, CER, PIP, and PS) across varying 5-HT concentrations and the average order parameter of both leaflets facing extracellular cytosolic sides. (B) Lipid order parameter of the membrane averaged over both leaflets. (C) Leaflet-wise individual order parameter. (D) Average Area Per Lipid (APL) computed independently for both leaflets. (E) Bilayer thickness across different 5-HT concentrations showing membrane thickening.

### 5-HT Induces Tighter Lateral Packing and Thickening in Heterogeneous Brain Plasma Membrane Unlike Single Component Membrane

While the previous analysis establishes that 5-HT actively modulates selective lipid order, the area per lipid (APL) quantifies the magnitude of the 5-HT induced structural alterations in the membrane. Initially, two systems are set up: a symmetric single-component POPC membrane, incorporating the trans-membrane Y4R-PP complex in the absence and presence of 6 mM 5-HT concentration, as reported in previous experiments^10^. Upon addition of 5-HT, the area per lipid shows a prominent increase from 0.66 *±* 0.002 nm^2^ to 0.68 *±* 0.001 nm^2^. This result indicates that the membrane becomes more fluidic in the presence of 5-HT (Figure S1 of the SI), which is in agreement with the experimental results^10^. Next, a comparative analysis between the pure brain plasma membrane from the literature^20^ and the protein-embedded control system from our simulation (S1) is conducted. The average area per lipid (APL) for the pure brain plasma membrane^20,26^ is 0.462 *±* 0.001 nm^2^ for the extracellular region and 0.497 *±* 0.001 nm^2^ for the cytosolic region^20^, whereas upon incorporation of the trans-membrane protein, the APL decreases to 0.414 *±* 0.002 nm^2^ for extracellular region and 0.474 *±* 0.002 nm^2^ for the cytosolic region. The incorporation of the trans-membrane protein decreases the average APL. Such a reduction indicates a tighter lateral packing of the lipid molecules around the protein core, which is consistent with the characteristic ordering effects typically exerted by transmembrane domains. However, upon addition of 5-HT, the APL does not increase as would be expected for a single-component membrane. Instead, the cytosolic APL decreases from 0 mM to 6 mM and remains nearly unchanged at 50 mM. The extracellular region exhibits a minor but progressive decrease from 0 −*>* 6− *>* 50 mM. These changes indicate a modest reduction in the lateral area occupied by individual lipids following 5-HT incorporation, with most of the adjustment occurring at 6 mM in the cytosolic region and showing a more gradual response in the extracellular region (Figure6 D). The APL changes are accompanied by a small increase in bilayer thickness, shown in figure 6. The thickness increases from S1-S3. The concurrent decrease in APL and increase in bilayer thickness are consistent with the order-parameter analysis, which shows a modest increase in both the overall and leaflet-resolved *S*_*CH*_ (Figure 6 B) values with increasing 5-HT concentration. These observations decipher that the membrane undergoes a subtle increase in lateral packing and bilayer compaction, unlike the membrane expansion and thinning reported for the single-component lipid systems. This behavior further highlights the importance of membrane composition or lipidomics in determining the physical response to 5-HT.

### 5-HT Alters Intramolecular Communication Pathways among Y4R Residues Experimentally Implicated in PP Recognition and Receptor Activation

To map the communication pathway within the Y4R-PP complex, the WISP analysis is employed. Combined study with double-cycle mutagenesis and three-dimensional model construction with available GPCR crystal structures reveals that PP recognition and Y4R activation involve multiple residues distributed across TM2, TM6, and TM7, including Tyr^2.64^, Asp^2.68^, Asn^6.55^, Asn^7.32^, and Phe^7.3521^. Among these residues, Tyr2.64 (referred to as TYR101 in our study) of TM2 is found to have the highest contact frequency with PP, as confirmed by GetContact^27^ analysis of our simulation trajectories as shown in the Figure S3 of the SI. TYR101 of Y4R is considered as the source. Asp^6.59^ of TM6, (referred to as ASP289 in our simulation), is selected as the sink because mutagenesis studies establish the functional importance of Asp^6.59^ in Y-receptor ligand binding and signal transduction^24,28^. Although these residues are not the only determinants of receptor function, their experimentally established roles provide a rationale for defining them as the endpoints of the WISP analysis. This approach allows us to investigate the receptor residues involved in communication between these two functionally important regions. Thus, TYR101 and ASP289 are defined as the signal source and sink, respectively.

Node degeneracy identifies the most frequently utilized residues across all optimal and suboptimal pathways.

Node degeneracy is the fraction of the 300 shortest paths that pass through a given residue, and it tracks the frequency of residue involvement in communication from the source (TYR101) to the sink (ASP289). Thus, high node degeneracy of a residue indicates its high contribution to the communication path. In the 5-HT-free state (S1), the communication propagates from the PP-ligand binding residue of Y4R (source, TYR101) to the sink, ASP289, through a highly coordinated pathway along the protein backbone. In this path, TRP107, PHE109, LEU113, LYS115, SER178, TRP205, LEU207, HSD210, and THR212 (shown in Figure 7A) have high node degeneracy and therefore act as primary contributing residues that physically stabilize the communication along Y4R protein. Among the residues identified in our calculations, TRP107 and CYS114 have previously been found relevant in Y4R function, although their roles differ. TRP107, located in the extracellular region of the receptor, has been experimentally associated with PP-induced receptor activity, as its substitution with alanine substantially reduces PP potency^21,28^. CYS114, in contrast, forms a disulfide bond with CYS201 that contributes to the structural stabilization of Y4R^29^. These findings provide experimental context for the residues identified in our communication pathways, while distinguishing the functional relevance of TRP107 from the structural role of CYS114. The residues identified for our calculations contributing to the communication path, corresponds to L2, L4, H3 and H5.

**Figure 7.**
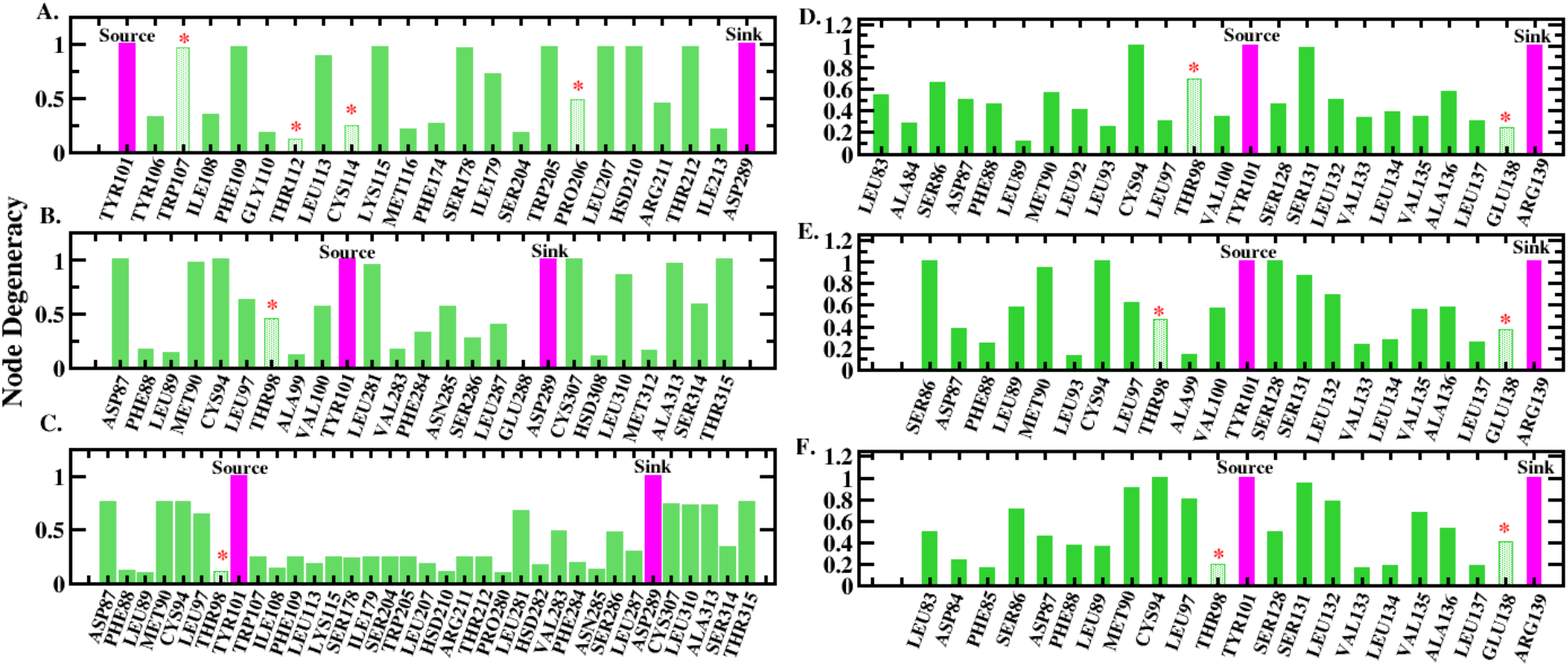
Node degeneracy profiles of residues involved in the WISP-derived communication pathways between TYR101 as the source and ASP289 as the sink at A) 0 mM (S1), B) 6 mM (S2), and C) 50 mM (S3) serotonin (5-HT). TYR101 as the source and ARG139 as the sink at D) 0 mM, E) 6 mM, and F) 50 mM 5-HT, respectively. Node degeneracy represents the fraction of the 300 shortest communication pathways that pass through a given residue. Magenta bars indicate the source and sink residues, while green bars represent other residues involved in the communication pathways. Asterisks indicate residues experimentally implicated in PP recognition and binding. The concentration-dependent changes in node degeneracy highlight the redistribution of residue contributions to intramolecular communication between the extracellular PP-binding region and either the PP binding-associated activation region, ASP289, or the intracellular G-protein-coupling associated residue, ARG139.

As the 5-HT concentration increases from 0−6 mM, a profound rewiring of the extracellular communication pathway is observed. The Y4R communicates through an alternative pathway involving ASP87, MET90, CYS94, LEU281, CYS307, LE310, ALA313, and THR315 (shown in Figure 7 B), with the highest node degeneracy. At 50 mM 5-HT (S3), the WISP-derived pathway retains essentially the same residues that are observed at 6 mM (S2). However, the two paths of 6 and 50 mM differ markedly in the degeneracy of the residues. At 50 mM, the node degeneracy of these same residues decreases, while a larger number of alternative residues acquire moderate degeneracy (shown in Figure 7 C). The corresponding pictorial representations of the optimal and suboptimal communication pathways are mapped directly onto the three-dimensional structure of the Y4R-PP complex for three concentrations and shown in Figure 8.

**Figure 8.**
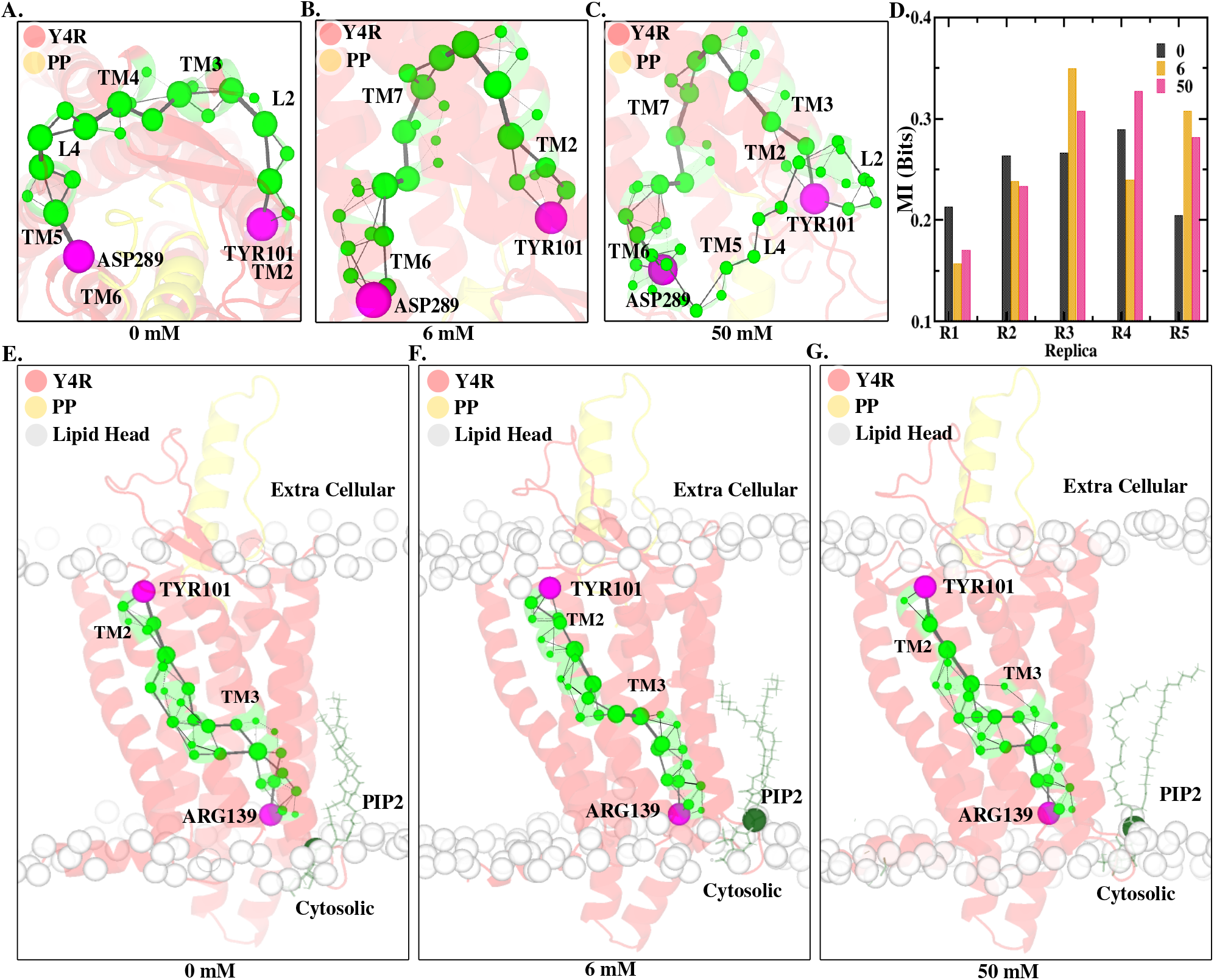
(A–C) Three-dimensional representations of the communication pathways from TYR101 to ASP289 in the Y4R–PP complex at (A) 0 mM (S1), (B) 6 mM (S2), and (C) 50 mM (S3) 5-HT. Spheres represent residues (nodes), with sphere size proportional to node degeneracy, which reflects the frequency with which a residue occurs among the computed communication pathways. Black lines represent residue–residue contacts (edges), with line thickness proportional to edge degeneracy; thicker edges indicate more frequently utilized communication routes. Green spheres represent residues involved in the communication pathways, while magenta spheres indicate the source (TYR101) and sink (ASP289). (D) Mutual information (MI) between the TYR101–LEU31 and TYR101–ASP289 distance series across five independent replicas at each 5-HT concentration. In the absence of 5-HT, MI exceeds 0.2 bits in all replicas, indicating statistical coupling between the PP-binding interface and the activation-associated region of Y4R. The changes in MI upon increasing 5-HT concentration suggest concentration-dependent modulation of this distance-based coupling. (E–G) Three-dimensional representations of the communication pathways from TYR101 to ARG139, located in the conserved E/DRY motif region of TM3, at (E) 0 mM, (F) 6 mM, and (G) 50 mM 5-HT. These pathways illustrate the redistribution of intramolecular communication from the extracellular PP-recognition region toward the intracellular G-protein-coupling interface. Lipid headgroups are shown as grey spheres, and PIP2, the closest lipids to Y4R, is indicated in dark green. Remaining lipids and water are not shown for the clarity.

### Serotonin-Dependent Modulation of Distance-Based Coupling Between the Y4R–PP Binding Interface and Activation-Associated Residue

The statistical coupling between the Y4R–PP binding interface, represented by the TYR101–LEU31 distance, and the receptor’s signaling-associated residue ASP289 is quantified using mutual information (MI), as described in Equation (12). Inter-residue distances (TYR101-ASP289 vs TYR101-LEU31) provide a structural readout of the dynamic relationship between ligand recognition and receptor communication. Changes in these distances (as shown in Figure S4-S5 in the SI) and their statistical dependence can indicate altered coupling between the PP-binding region and the receptor’s intracellular signaling machinery. In all replicas, the TYR101-ASP289 and TYR101-LEU31 distances fluctuate throughout the 20 ns trajectories and the fluctuations appear to be correlated. This temporal correlation, quantified by the autocorrelation function of each distance series, is used to set the block length of the permutation. In the absence of 5-HT, MI values exceeding 0.2 bits across all independent replicas indicate measurable statistical coupling between the binding-interface distance and the ASP289-associated distance (Figure7D) . As the 5-HT concentration increases from 0 to 6 and 50 mM, the MI values change across replicas, suggesting that serotonin concentration modulates the strength of this distance-based coupling. The significance of this coupling is assessed with a block-permutation test. At all three serotonin concentrations, the pooled MI lies well outside the corresponding null distribution (*p* = 0.002 for 0, 6 and 50 mM; which is shown in S7 of SI), and the MI of every individual replica is far above its null mean, indicating statistically significant coupling in all cases. Here *p* = 0.002 is the smallest value attainable with 500 permutations, i.e. no permuted data set reached the observed MI. From a chemical neuroscience perspective, these findings support a model in which serotonin, without requiring direct receptor contact, may influence how PP-binding-associated structural fluctuations are communicated toward a signaling-associated region of Y4R. Importantly, MI quantifies statistical dependence rather than causal information transfer; therefore, these results indicate concentration-dependent reorganization of intra-receptor communication, not direct proof of altered receptor activation or downstream signaling.

### 5-HT Alters Communication Pathways between Literature-Identified Y4R Residues Related to PP Binding and G-Protein Coupling, Located in Opposing Membrane Leaflets

To further investigate intramolecular communication within Y4R connecting PP recognition and Y4R activation to G-protein binding and activation, ARG139 (Arg^3.50^), located within the conserved E/DRY motif region of transmembrane helix 3 (TM3), is considered as an alternative sink. The E/DRY motif is a key regulatory element in class A G-protein-coupled receptors (GPCRs), contributing to the stabilization and interconversion of receptor conformational states associated with activation and G-protein coupling^30^. Using ARG139 as the sink allows us to examine communication pathways extending from the extracellular TYR101 (Tyr^2.64^) source towards the intracellular G-protein-coupling interface. As the G protein is absent from the simulated systems, this analysis characterizes intrareceptor communication towards the coupling interface rather than direct receptor–G-protein interactions. Across the three 5-HT concentrations (0, 6, and 50 mM; 7D–F), several residues recur in the computed source-to-sink pathways, suggesting that the pathways share common structural elements while the contribution of individual residues varies with serotonin concentration. The residues in the communication paths are part of helices 2 and 3 of Y4R. Notably, THR98 (Thr^2.61^), which has been implicated in the PP-binding region and in the activity of the positive allosteric modulation, exhibits a progressive reduction in node degeneracy as the 5-HT concentration increases. Previous structural and mutagenesis studies support a role for Thr^2.61^ (THR98) in PP-related interactions^31^. The reduced node degeneracy of THR98 indicates that it occurs less frequently among the selected communication pathways at higher serotonin concentrations. Together, these findings suggest that serotonin concentration is associated with changes in the intramolecular communication between the extracellular ligand-recognition region and the intracellular face of Y4R.

The observed changes in intramolecular communication from the extracellular ligand-recognition region towards the cytosolic face of Y4R may have implications for receptor–membrane interactions, particularly given the presence of PIP2 in the cytosolic leaflet. This raises the possibility that 5HT-induced changes in receptor communication are associated with the local lipid environment at the intracellular interface, although a direct mechanistic link between PIP2 and the observed pathways remains to be established. Note, changes in node degeneracy reflect the frequency with which a residue occurs among the computed pathways and do not, by themselves, establish changes in receptor activation or G-protein signalling efficacy.

### 5-HT Modulates Receptor Fluctuations and Orientations of Regions Involved in Communication Pathways for PP Binding and Y4R Activation

To monitor whether these 5-HT concentration influences receptor residue-wise fluctuations, root-mean-square fluctuation (RMSF) analysis are done for all three concentrations. In the presence of 5-HT, there is an overall reduction in receptor fluctuations, specifically in the linker L2, L4 and H5, indicating reduced local conformational flexibility (Figure A of S8 in SI panel 1). Interestingly, in the absence of 5-HT, the dominant residues that contribute to the communication pathway are primarily associated with L2, L4, H3 and H5 when ASP289 has been considered as a sink and H2 and H3, when ARG139 has been considered as a sink. Most of which have reduced RMSF upon incorporation of 5-HT. The reduction in fluctuations in L2 and L4, is demonstrated in the porcupine diagram shown in Figure B of S8 in the SI, which depicts the directions and relative magnitudes of residue motions. These changes suggest that 5-HT modulates the local conformational flexibility of selective regions of the receptor rather than inducing a uniform global change. The tilt-angle analysis further reveals changes in the orientation of the selective linkers, L2, L4, and helices, H2, H3, H4 and H5 of Y4R with respect to the bilayer normal. With increasing 5-HT concentration, the tilt of L2 decreases progressively, whereas the same for L4 increases. Among the transmembrane helices, H2 shows an increase in tilt from 0 to 6 mM followed by a decrease at 50 mM, although its tilt remains higher than that of the control, while the tilt of H3 and H6 decreases with increasing 5-HT concentration (Figure C of S8 in SI panel 2 and 3). These selective changes in orientation, together with the overall reduction in RMSF, indicate that 5-HT modifies the local fluctuations and orientations of Y4R while preserving the overall transmembrane architecture. As mentioned, the regions showing these changes correspond to the redistribution of the receptor communication pathways. In the absence of 5-HT, the dominant residues are primarily associated with L2, L4, H3, and H5, whereas incorporation of 5-HT shifts the residues toward H2, H6, and H7 for ASP289 but the communication pathways are conserverd in H2 and H3 for ARG139 at all 5-HT concnetrations. Thus, the specific linkers and helices that exhibit reduced fluctuations and altered orientations upon 5-HT binding also undergo redistribution of their intramolecular communication pathways, linking local structural changes to altered receptor communication.

## Conclusion

Understanding serotonergic modulation of Y4R signaling and PP recognition is important for elucidating gut–brain communication and neuropsychiatric function. As a Class A GPCR, Y4R function is shaped not only by PP binding, but also by the surrounding membrane environment. Thus, brain plasma membrane heterogeneity may influence receptor conformation, ligand recognition, and signaling, linking membrane composition to GPCR-mediated neuronal function. Earlier experiments demonstrate that neurotransmitter, 5-HT, modulates the binding kinetics of PP to its cognate Y4R receptor without directly interacting with the receptor, indicating a membrane-mediated allosteric mechanism and highlighting the membrane as a potential target for pharmacological modulation^10^. However, the molecular pathways that are involved in 5-HT induced Y4R communication for its PP recognition and activation remains unknown. Moreover, the role of lipid diversity of the heterogeneous brain plasma membrane on 5-HT altered Y4R-PP communication remains unresolved. To understand the molecular mechanism of 5-HT-induced alterations in the noncognate Y4R-PP complex, a total of 825.9 ns AA MD simulations are conducted for three distinct concentrations of 0, 6, 50 mM 5-HT using the CHARMM36m force field at 310 K. The amphiphilic nature of 5-HT allows it to preferentially localize at the lipid–water interface, positioning itself close to the lipid headgroup region throughout all concentrations. 5-HT resides closer to the lipid heads, specifically, PC lipids, than the Y4R-PP signaling residues, confirming it as a non-cognate ligand to Y4R. However, the preferential contacts of 5-HT and the receptor with the zwitterionic lipids are mutually exclusive. 5-HT partitions into the distinct zwitterionic PC lipid microenvironment. The lipid preference of the receptor remains conserved across all 5-HT concentrations. The anionic PIP2 exhibits the strongest enrichment within the receptor contact shell, followed by PS, PE, SM, CER, CHOL, and POPC. The close agreement between the D–E index and protein–lipid contact frequency, together with the observation that the anionic lipid PIP2 contacts the receptor more frequently than 5-HT, suggests that PIP2 becomes the dominant interaction partner within the receptor–5-HT microenvironment. Although at 6 mM, the anionic lipid, PIP2, has preferential contact with the receptor over 5-HT, it emerges as the dominant, mutually preferred species within both the receptor and 5-HT microenvironments, as 5-HT increases from 6 to 50 mM. Thus, PIP2 executes a biphasic sorting for the receptor and 5-HT, despite the low abundance of PIP2 in the brain plasma membrane. This concentration-dependent redistribution is accompanied by selective remodeling of the lipid–lipid interaction network, particularly within the anionic PIP2-PS-associated network, while the stressed lipid (CER-SM) associated network remains comparatively conserved. CER and SM are important bio-active sphingolipid components implicated in membrane organization and cellular stress-related signaling^25,32,33^. The persistence of the stress-associated lipid network across 5-HT concentrations suggests that this connectivity represents an intrinsic structural feature of the brain plasma membrane rather than a direct effect of 5-HT. Therefore, 5-HT selectively remodels anionic lipid-associated networks within the heterogeneous brain plasma membrane. The membrane-level reorganization is further reflected in its physical properties. Unlike previous studies of single-component POPC membranes reporting 5-HT-induced membrane disordering^9,10^, the heterogeneous brain plasma membrane exhibits a modest increase in lipid-chain order together with a reduction in area per lipid and a slight increase in bilayer thickness with increasing 5-HT concentration. These changes indicate subtle membrane compaction rather than global membrane disordering. The contrasting behavior emphasizes the importance of lipid diversity and membrane organization in determining the biophysical response to 5-HT.

To quantify the effect of 5-HT on Y4R communication, relevant for serotonergic signaling of Y4R, WISP analysis is performed between two residues of Y4R, TYR101 and ASP289, experimentally known PP binding and activation residue of Y4R, respectively^21,34^. In the absence of 5-HT, the communication between TYR101 and ASP289 is predominantly mediated through TRP107, PHE109, LEU113, LYS115, SER178, TRP205, LEU207, HSD210, and THR212, defining the principal communication route of the native receptor. The TRP107 residue identified in our calculations corresponds to an experimentally characterized^21^ Y4R residue whose substitution with alanine results in a 107-fold reduction in PP potency, highlighting its importance in PP-bound receptor activation. Upon addition of 5-HT at 6 mM, the native communication route is substantially changed, where TRP107 is no longer the communicating residue. Instead, ASP87, MET90, CYS94, LEU281, CYS307, LE310, ALA313, and THR315 emerge as the principal high-degeneracy residues. With a further increase in 5-HT concentration to 50 mM, the overall communication route of 6 mM is preserved with a significant reduction in the node degeneracy of the dominant residues, while a larger number of alternative residues acquire moderate degeneracy. This indicates that increasing 5-HT concentration progressively weakens the dominant communication hubs and recruits alternative residue pathways, and thereby providing a molecular basis for concentration-dependent serotonergic modulation of Y4R activation. To further examine communication towards the intracellular G-protein-coupling interface, ARG139 (Arg^3.50^), located within the conserved E/DRY motif^30,35^, is considered an alternative sink. The resulting pathways reveal serotonin-dependent changes in intramolecular communication from the extracellular TYR101 source towards the cytosolic face of Y4R. Given the enrichment of PIP2 in the receptor’s cytosolic contact environment, these changes may be associated with the local lipid environment surrounding the intracellular interface, although a direct mechanistic link remains to be established. The linkers and helices involved in communication paths between TYR101 and ARG139 or ASP289 have reduced fluctuations and alterations in tilt with respect to the bilayer normal, upon increasing 5-HT concentrations.

Taken together, our simulations reveal a membrane-mediated molecular mechanism through which 5-HT can modulate Y4R activation in the absence of a direct Y4R–5-HT contact. Our results confirm the experimental prediction of membrane allostericity, in which lipid reorganization provides a route for communication pathways rather than directly demonstrating receptor activation or downstream signaling relevant for serotonergic signal transduction. More broadly, this work provides an atomistic framework for understanding how membrane composition and lipid-network organization contribute to GPCR regulation and changes the optimal communication pathway, offers a basis for exploring membrane-mediated allosteric behavior in neuronal GPCR pharmacology. These findings open compelling new avenues for both pharmacological research and advanced computational biophysics. Since 5-HT alters receptor signaling via the lipid matrix rather than direct orthosteric binding, future drug discovery efforts could pivot toward lipid-targeted therapeutics as an alternative to protein.

## Method

### Quantum Chemical Calculation

The equilibrium geometry of the aliphatic terminal amine tautomer of protonated serotonin^36^ is obtained from the MP2/cc-pVDZ-optimized structure reported by Omidyan et. al.^37^. Since MP2/cc-pVDZ level of theory explicitly incorporates electron correlation and provides a high-quality equilibrium geometry for this protonated small-molecule cation, the optimized geometry is adopted without further re-optimization. Using this geometry without modification, a single-point calculation is performed at the B3LYP/6-311++G(d,p) level of theory in Gaussian-09^38^ to compute atomic partial charges using the CHELPG (Charges from Electrostatic Potentials using a Grid-based method) scheme. The resulting CHELPG charges are assigned to the corresponding atoms during force-field parameterization in the CHARMM General Force Field (CGenFF) server^39^, rather than allowing CGenFF to assign its own default charges via empirical bond-charge increment rules. This step ensures that the atomic partial charges used for AA MD simulations are derived from a quantum-mechanically consistent electrostatic potential. All remaining force field parameters (bonded and Lennard-Jones terms) are obtained from CGenFF.

### System Setup

The initial structure of the active-state human neuropeptide Y4 receptor (Y_4_R) in complex with its endogenous ligand, pancreatic polypeptide (PP), is obtained from the cryo-EM coordinates (PDB ID: 7X9C)^22^. The receptor–ligand complex is isolated from the heterotrimeric G protein present in the experimental structure. Residues 1–34 of Y4R are truncated, and the protein spans residues Gln35 to Gln341. Missing internal residues within the resolved protein construct are reconstructed using the SWISS-MODEL^40^ homology modeling server. The final simulation model comprises a core bundle of seven transmembrane helices (TM1–TM7) that traverse the hydrophobic lipid bilayer, connecting the extracellular environment to the cytosolic side. Unstructured terminal regions—specifically the N-terminus preceding Gln35 and the C-terminus beyond Gln341 are truncated and excluded from the system.

The receptor-ligand transmembrane Y4R-PP complex is embedded into this asymmetric plasma membrane model using the CHARMM-GUI Membrane Builder^41^.

The protein-ligand embedded membrane is then solvated with 36998 TIP4P/2005^42^ water molecules and neutralized with 44 KCl. To systematically investigate the concentration-dependent effects of the 5-HT, three distinct bilayers are referred to as: S1 (0 mM, control), S2 (6 mM), S3 (50 mM) (see Table 1). 6 mM concentration is chosen to compare our results with the previous experiment that shows the 5-HT acts as an allosteric modulator at this concentration^10^. To capture the concentration-dependent effects of 5-HT on the protein-ligand, 50 mM is included. This approach allows us to systematically examine how varying 5-HT concentrations influence the properties of the brain plasma membrane as well as the receptor and the ligand.

**Table 1.** Summary of simulated systems of the model brain plasma membrane (BPM) with a total simulation time of 825.9 ns.

| **System** | **Composition** | **5-HT (mM)** | **No. 5-HT** | **No. Cl<sup>−</sup>** | **Box (nm)** |  |  | **NPT** (total for 5 replicas) (ns) | **NVT** (total for 5 replicas) (ns) |
|---|---|---|---|---|---|---|---|---|---|
|  |  |  |  |  | x | y | z |  |  |
| S1 | BPM + Protein | 0 | 0 | 0 | 10.75 | 10.75 | 10.30 | 420.2 | 100.1 |
| S2 | BPM + Protein + 5-HT | 6 | 7 | 7 | 10.66 | 10.66 | 10.36 | 45.2 | 100.1 |
| S3 | BPM + Protein + 5-HT | 50 | 57 | 57 | 10.69 | 10.69 | 10.26 | 60.20 | 100.1 |

### Simulation Details

AA MD simulations are performed using the CHARMM36m force field^43^. The control system (S1) is energy minimized utilizing the steepest descent algorithm^44^ to eliminate potential steric clashes. Subsequently, a multi-step restrained equilibration protocol is executed. Initially, a 100 ps simulation is performed in the NVT ensemble with a 2 fs time step. During this step, position restraints of 4000, 2000, and 1000 kJ/mol/nm^2^ are applied to the protein backbone, side chains, and dihedral angles, respectively, along a 1000 kJ/mol/nm^2^ restraint on the lipid molecules. The restraint strengths of the backbone, sidechain, dihedrals, of the protein and lipid heads are subsequently reduced to 2000, 1000, 400, and 400 kJ/mol/nm^2^, respectively, to run an additional 100 ps NVT simulation. The S1 system is then transitioned to the NPT ensemble. Three successive NPT stages of 100 ps each (totaling 300 ps) are performed, with restraints progressively reduced to 1000, 500, 200, and 400 kJ/mol/nm^2^ in the first stage; 500, 200, 200, and 200 kJ/mol/nm^2^ in the second; and 200, 50, 100, and 40 kJ/mol/nm^2^ in the third for the protein and lipids respectively. Following these restrained steps, all position restraints on both the protein and the lipid molecules are completely removed. A prolonged 415.2 ns unrestrained NPT simulation is then performed to fully relax the system. Subsequent NPT equilibrations are performed until the area per lipid of the bilayer converges and attains equilibrium (data provided in Figure S1 of the supporting information (SI)). The final equilibrated configuration of the trajectory is used as the initial structure of the 5 replicas generated for S1 as well as for the remaining bilayers, S2 and S3. The 5-HT molecules are randomly incorporated into the extracellular and cytosolic water of S2 and S3 to attain the desired concentration of 5-HT. Each of these newly formed 5-HT-containing bilayers is then independently energy-minimized using the steepest-descent algorithm^44^. Subsequent 40.2, 55.2 ns NPT equilibrations are performed for S2 and S3 respectively, until the area per lipid of the bilayer converges and attains equilibrium. The final configurations of equilibrated S2 and S3 are used as the initial configurations of all 5 replicas by reassigning initial atomic velocities from a Maxwell–Boltzmann distribution at 310 K using unique random seeds. Each replica is then subjected to a 1 ns unrestrained NPT equilibration to verify system stability, followed by a 20 ns NVT production trajectory, with frames saved every 20 ps. The details of the total run lengths are mentioned in Table 1. Throughout all the NPT equilibration, a constant pressure of 1 bar is maintained using the semi-isotropic Berendsen pressure coupling method^45^ with a coupling constant of 1.0 ps. Non-bonded electrostatic and van der Waals interactions are truncated at a distance of 1.2 nm, and long-range electrostatic interactions are computed utilizing the Particle Mesh Ewald (PME) method^46^ with a grid spacing of 0.16 nm. Periodic boundary conditions are applied in all three spatial dimensions, and all the hydrogen bonds are constrained using the LINCS algorithm^47^ with an order of 4. The temperature is kept constant at 310 K utilizing the velocity-rescaling thermostat^48^ with a coupling constant of 1.0 ps throughout both the equilibration and production runs. All the simulations are performed using Gromacs(2022.5)^49^.

## Analysis

All analyses are performed using the production run trajectories of five independent replicas for each system (S1-S3). For visualization, the final configuration from one representative replica of each system is shown in Figure2.

### Radial Distribution Function

The three-dimensional radial distribution function, *g*(*r*), quantifies the radial association of neighboring molecules around each other and is calculated by the following equation:

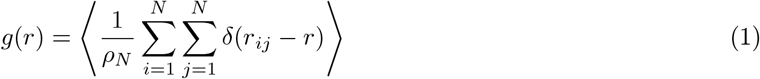

where *N* denotes the total number of molecules, *ρ*_*N*_ the density of the molecules, *r*_*ij*_ the distance between particle *i* and *j*, and ⟨…⟩ represents the ensemble average.

All final *g*(*r*) profiles represent the ensemble average calculated across the five independent replicas for each respective system.

### Depletion Enrichment Index (D–E index)

To characterize the local lipid environment surrounding the Y4R receptor, the lipid depletion–enrichment (DE) index is calculated following the approach of Corradi et al.^50^, using a 0.65 nm cutoff between all atoms of the protein and all atoms of each lipid species to define a lipid molecule belonging to the contact shell of the protein. For each lipid type, the DE index is calculated as the ratio of its fractional abundance within the contact shell to its fractional abundance in the bulk membrane as follows,

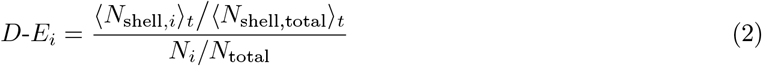

where *N*_shell,*i*_ is the number of lipid molecules of type *i, N*_*i*_ is the total number of molecule of lipid type *i*. ⟨*N*_shell,*i*_ ⟩_*t*_ is the time-averaged number of lipid molecules of type *i* found within the protein’s contact shell, and ⟨*N*_shell, total_⟩ _*t*_ is the time-averaged total number of lipid molecules of all types combined found within that same shell, such that ⟨*N*_shell,*i t*_*/ N*_shell, total_⟩ _*t*_ gives the fractional abundance of lipid *i* within the shell. Similarly, *N*_*i*_ is the total number of molecules of lipid type *i* in the entire membrane, and *N*_total_ is the total number of lipid molecules of all types in the entire membrane, such that *N*_*i*_*/N*_total_ gives the fractional abundance of lipid *i* in the bulk membrane. The DE index (*D*-*E*_*i*_) is therefore the ratio of the shell fractional abundance to the bulk fractional abundance. It indicates whether lipid *i* makes up a larger or smaller fraction of the protein’s shell than it does of the membrane as a whole. Values of D − E_*i*_ *>* 1 indicate enrichment, whereas values of D − E_*i*_ *<* 1 indicate depletion relative to the bulk composition, with D − E_*i*_ = 1 indicating bulk-like, non-preferential association. All calculations are performed independently for each replica and averaged across the five replicas, with the standard deviation reported as the associated uncertainty.

### Contact frequency

Protein-lipid interactions are quantified using a distance cutoff of 0.65 nm applied between all atoms of the protein and all atoms of each lipid species *L*. This cutoff corresponds to the outer boundary of the second hydration shell around heavy atoms, thereby capturing both direct non-covalent contacts and short-range interactions between the interacting species. At each simulation frame *t*, a lipid molecule is considered to be in contact with the protein if any of its atoms lies within 0.65 nm of any protein atom. The number of distinct contacting lipid molecules of species *L* at frame *t*, denoted as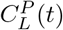, is normalized by the total number of molecules of that lipid species present in the system (*N*_*L*_). The normalized protein–lipid contact fraction (*CF*_*P* −*L*_) is then obtained by time-averaging over the total number of trajectory frames (*N*_*f*_ ):

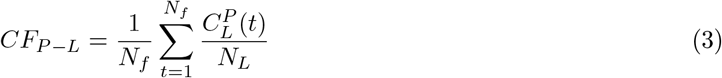

where *CF*_*P* −*L*_ represents the time-averaged fraction of lipid species *L* in contact with the protein. Similarly, 5-HT–lipid contacts are quantified using the same 0.65 nm cutoff applied between all atoms of 5-HT and all atoms of each lipid species *L*. The number of distinct lipid molecules of species *L* contacting 5-HT at frame *t*, 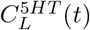, is normalized by both the total number of 5-HT molecules (*N*_5*HT*_ ) and the total number of molecules of lipid species *L* (*N*_*L*_). Time-averaging over all trajectory frames yields the normalized 5-HT–lipid contact fraction (*CF*_5*HT* −*L*_):

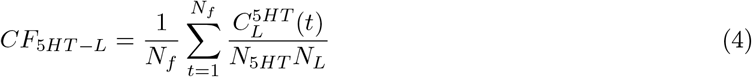

For both analyses, calculations are performed independently for each of the five replicas, and the resulting values are reported as the mean *±* standard deviation across the replicas. The depletion–enrichment (*D*-*E*) index and contact fraction analyses are implemented using the MDAnalysis package^51^ in Python, supported by NumPy^52^, pandas^53^, and seaborn^54^ libraries^51,55^.

### Lipid-Lipid Enrichment and Network analysis

Building upon these contact analyses, lipid–lipid co-localization is examined by constructing an interaction network using the NetworkX Python package^56^, in which each lipid species is represented as a node. Unlike the DE index described above, which evaluates the abundance of each lipid species within the contact shell of a single external reference point receptor, this analysis evaluates the joint occurrence of two lipid species drawn from the same membrane pool and is therefore defined for every pair of lipid species rather than for each species individually.For each replica, pairwise lipid–lipid contacts are identified using a 0.65 nm cutoff applied to the center-of-mass (COM) distance between lipid molecules. The contacts are averaged over the simulation trajectory to obtain the mean number of contacts per frame between each pair of lipid species, ⟨*n*_*ij*_⟩. This quantity is compared with the number of contacts expected under a null model of random, non-preferential lipid mixing, based on the relative abundance of each lipid species in the system, to obtain a pairwise enrichment ratio:

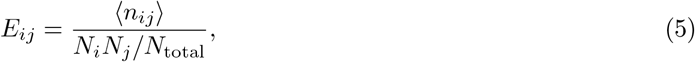

where *N*_*i*_ and *N*_*j*_ are the total numbers of molecules of lipid species *i* and *j* in the system, respectively, and *N*_total_ is the total number of lipid molecules. Values of *E*_*ij*_ *>* 1 indicate preferential co-localization between lipid species *i* and *j* relative to random mixing, whereas *E*_*ij*_ *<* 1 indicates reduced co-localization or avoidance. Calculations are performed independently for each of the five replicas and subsequently averaged, with the standard deviation across replicas reported as the associated uncertainty. In the resulting network representation, an edge is drawn between two lipid nodes only when their mean enrichment ratio exceeds the threshold *E*_*ij*_ *>* 1.2. The edge color and width are scaled according to *E*_*ij*_ to highlight the strongest lipid-lipid co-localization pairs.

### Order Parameter (*S*_*CH*_)

To elucidate the structural organization of the lipid bilayer, the acyl-chain order parameter, *S*_*CH*_, is calculated across all replicas for individual lipids within each leaflet using the `g_order` analysis toolFirst, lipid molecules are identified and classified into distinct lipid types based on their atomic sequences, residue identities, and bonding connectivity. Subsequently, user-defined selections of heavy atoms and hydrogens are used to identify the corresponding bonds for calculating segmental order parameters. For systems simulated with explicit hydrogen representations, the order parameter is mathematically defined as

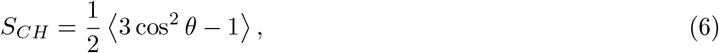

where *θ* represents the instantaneous angle between the carbon-hydrogen (C-H) bond vector and the normal vector to the local lipid bilayer. The angular brackets, ⟨*· · ·* ⟩, denote both the temporal average over the simulation trajectory and the molecular ensemble average across all equivalent lipid instances.In accordance with standard nuclear magnetic resonance (NMR) experimental conventions, the output values are reported as the sign-inverted magnitude, −*S*_*CH*_ . Consequently, higher values of −*S*_*CH*_ correspond to restricted motional freedom and a more ordered membrane state, whereas lower values indicate increased conformational flexibility and structural disorder induced by the intercalated 5-HT.

### Area per Lipid and Bilayer Thickness

Furthermore, the macroscopic physical properties of the membrane, specifically the bilayer thickness and the area per lipid (APL), are computed using the LiPyphilic package^57^. This object-oriented Python toolkit is explicitly designed for the analysis of lipid bilayer simulations and is built directly upon the MDAnalysis framework^51^ alongside other scientific Python packages such as SciPy, Numpy^52,58^.

### Weighted Implementation of Suboptimal Paths (WISP)

WISP uncovers the allosteric communication pathways between remote parts of a protein. To model how the communication propagates from the ligand-contacting region to the functional site of Y4R through a specific pathway of the protein, source and sink nodes are defined^59^. The source and the sink are designated as the protein residues that are experimentally known to be involved in PP ligand recognition and Y4R activation. Since there can be multiple residues involved in PP recognition and Y4R activation, the common residue that is experimentally known for PP recognition and that has the highest contact frequency with the ligand from our simulation is considered to be the source. Experimentally, it is known that TYR101 of Y4*R* is a contact hub with the LEU31 residue of ligand PP^21,34^. The same residue is found to have the highest contact with PP from our simulation. Thus, TYR101 of Y4*R* is considered as the source. The sink represents the key residue that defines the functional site of the receptor protein.

Since ASP289 is experimentally known to be crucial for PP recognition and Y4R activation^24^, it is considered the sink. The WISP analysis between TYR101 and ASP289 provides the communication path between two residues of Y4R involved in PP-recognition, binding, and Y4R activation. ARG139^3.50^ (ARG139 ) is selected as another candidate for the sink residue for investigating Y4R-mediated G-protein coupling because the Y4R–PP–Gi1 cryo-EM structure (PDB 7X9C) identifies direct contacts between this residue and the C-terminal *α*5 helix of G*α*_i1_. Furthermore, Arg^3.50^ occupies the conserved DRY motif of class A GPCRs, which plays a central role in receptor activation and intracellular G-protein engagement^21,35,60^. ARG139 and ASP 289 of Y4*R* are considered as activation-relevant residue for signal transduction^24,28,31^. To this end, we consider two sets of source and sink: TYR101-ASP289 and TYR101-ARG139. To characterize the dynamic network between the source and the sink, the primary communicating path (optimal path) as well as the slightly longer suboptimal paths are analyzed employing the Weighted Implementation of Suboptimal Paths (WISP)^61^ as implemented in the Bio3D package^62^. Five independent replicas are PBC corrected and concatenated. The protein is modeled as a network in which each residue is a node represented by its C*α* atom and an edge connects two residues that are in contact. Each edge is weighted by the correlation of the motions of the two residues it connects. The PP ligand is not included as a node. To represent the topology of the network, the Dynamic Cross-Correlation Matrix (DCCM) and the contact map are generated as the primary inputs for WISP, conducted via the Bio3D package.^62^. The dynamic cross-correlation matrix (DCCM, *C*_*ij*_) is calculated from C*α* fluctuations using the dccm function of Bio3D.

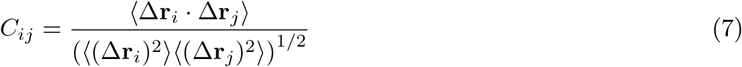

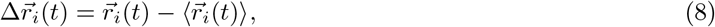

where, *N* represents the total number of nodes, with *i* and *j* denoting individual node indices. Additionally, Δ**r**_*i*_ and Δ**r**_*j*_ represent the displacement vectors of atoms *i* and *j* from their respective mean positions over the simulation trajectory. To ensure that strongly correlated residue pairs are close in network space, the correlation values are transformed into network edge lengths (*w*_*ij*_) using a negative logarithmic transformation, expressed as:

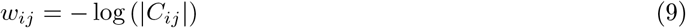

Under this transformation, strong correlation or anticorrelation (|*C*_*ij*_| →1) results in an edge length close to 0 (−log(1) = 0), indicating that the residues are close in network space. Conversely, weak or absent correlation (| *C*_*ij*_ |→0) yields a large edge length (−log(0) → ∞), meaning the residues are positioned far apart in the network.

A binary structural contact map is generated utilizing a 0.65 nm distance cutoff between any C*α* atom of the protein residue pairs. This specific threshold is selected to comprehensively capture dominant first-shell non-covalent interactions (such as van der Waals and electrostatic contacts) while excluding distant, noninteracting residue pairs. Concurrently, edges are retained only for residue pairs that are in contact and satisfy |*C*_*ij*_| ≥0.25, which effectively filters out weak, transient fluctuations arising from random thermal noise, ensuring the resulting pathways represent statistically robust, coupled motions, removes weakly correlated pairs. WISP computes shortest communication pathways bridging the spatially distant nodes of the Y4R receptor. Although our initial selection of 500 shortest paths between TYR101 and ASP289 was arbitrary, the minimum number of paths required to reliably predict node degeneracy is examined. By analyzing the trajectory across varying path counts, the degeneracy of key residues involved in the signal transduction is tracked. These results demonstrate that node degeneracy largely converges at 350 paths. The node degeneracy is calculated to determine the functional importance of specific residues by demonstrating how frequently they appear across the computed signaling pathways. It is defined as the number of paths that pass through residue *i*:

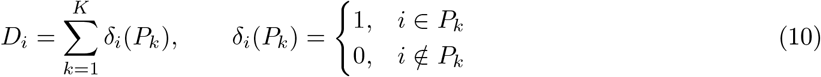

where *P*_*k*_ is the *k*-th shortest path and *K* is the total number of paths (*K* = 300). We divided by *K* to get the fraction of paths that use the residue:

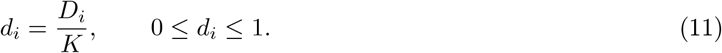

A residue with *d*_*i*_ close to 1 is used by almost all pathways and is likely important for signal transmission. A residue with *d*_*i*_ close to 0 is used by very few pathways.

### Mutual Information Analysis

Mutual information (MI) is calculated between the intramolecular distance *x*(*t*), defined between the C_*α*_ atoms of Y4R residues Tyr101 and Asp289, and the intermolecular distance *y*(*t*), defined between TYR101 of Y4R and LEU31 of PP. The analysis included 5, 000 frames from five independent replicas (1, 000 frames per replica). MI was calculated as

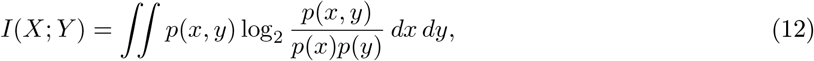

where *p*(*x, y*) is the joint probability density and *p*(*x*) and *p*(*y*) are the marginal probability densities of the two distances. MI quantifies their statistical dependence but does not establish causality or directionality.

Statistical significance is assessed using 500 block-permutation tests. The *x*(*t*) series is divided into contiguous blocks, which are randomly reordered within each replica while the corresponding *y*(*t*) series remains unchanged. Block length is determined from the autocorrelation decay times of both distance series, with a minimum of 5 frames and a maximum of 10% of each replica’s trajectory length. For each permutation, the shuffled data from all five replicas are pooled to calculate the null MI. The permutation *P*- value is calculated as

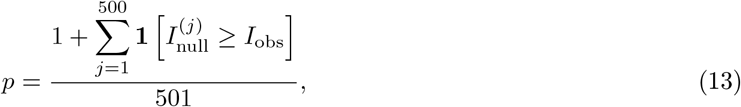

where *I*_obs_ is the observed pooled MI, 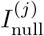 is the pooled MI from permutation *j*, and **1**[*·*] equals 1 when the stated condition is satisfied and 0 otherwise.

### Root Mean Square Fluctuation (RMSF) and Tilt Angle

To assess the local conformational flexibility of Y4R in response to changes in 5-HT concentration, the perresidue root mean square fluctuation (RMSF) of the receptor backbone is calculated using the ‘gmx rmsf’^49^ module of GROMACS. The RMSF is calculated from the positions of the protein backbone atoms according to

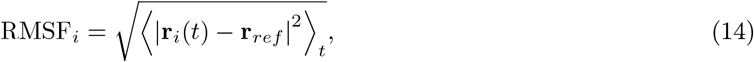

where **r**_*i*_(*t*) represents the position of the backbone atoms of residue *i* at time *t*. The initial structure of Y4R from the cryo-EM coordinates is used as the reference structure for the fitting. The RMSF therefore measures the extent to which the backbone atoms of each residue fluctuate around the respective experimental positions over the course of the simulation. Before calculating RMSF, the trajectories are least-square fitted to the experimental protein backbone to remove overall translational and rotational motion of the receptor. RMSF values are subsequently calculated on a residue-wise basis from the backbone atoms. The calculation is performed independently for each of the five replicas at 0, 6, and 50 mM 5-HT concentrations. For each residue, the RMSF values from the five independent replicas are averaged, and the standard deviation across replicas is reported as the associated uncertainty.

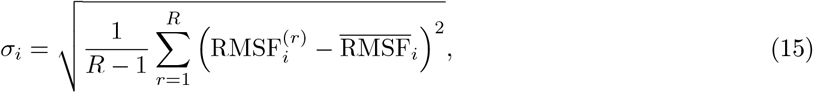

where *R* = 5 is the number of independent replicas and 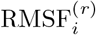 represents the RMSF of residue *i* in replica *r*.

The orientation of each transmembrane (TM) helix and the linkers (L) relative to the membrane is quantified by calculating its tilt angle with respect to the bilayer normal. For each TM helix and linkers the principal axis vector **â** are determined using principal component analysis (PCA) of the coordinates of its constituent C_*α*_ atoms. The coordinates 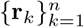 of the *n* residues comprising the helix are first mean-centered as

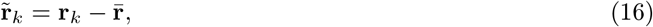

Where

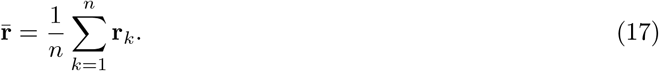

The 3 *×* 3 covariance matrix of the centered coordinates are then constructed as

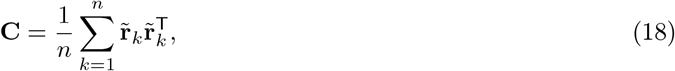

and diagonalized. The eigenvector corresponding to the largest eigenvalue of **C** are taken as the principal (long) axis of the helix, representing the best-fit straight-line direction through the helix backbone. This PCA-based definition is robust to helical curvature, in contrast to a simple vector connecting the N- and C-terminal residues, which may be distorted by bending along the helix.

The bilayer normal 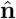 is taken as the unit vector along the *z*-axis, consistent with the membrane orientation in the simulation setup, in which the bilayer lies in the *xy*-plane. The tilt angle *θ* between the helix axis and the bilayer normal is calculated as

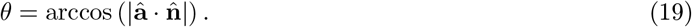

The absolute value of the dot product is used because the PCA-derived principal axis **â** has no intrinsic sign or directionality, thereby restricting *θ* to the range [0^*°*^, 90^*°*^]. A tilt angle of *θ* = 0^*°*^ corresponds to a helix oriented perpendicular to the membrane plane, i.e., fully spanning the bilayer with no lateral lean, whereas *θ* = 90^*°*^ corresponds to a helix oriented parallel to the membrane plane, i.e., lying approximately within the membrane plane. Tilt angles are calculated on a per-frame basis for each TM helix in each of the five replicas and reported as the mean *±* standard deviation across replicas. Visual inspection of the molecular trajectories and computed pathways is conducted using Visual Molecular Dynamics (VMD) and PyMOL.

## Supporting information

Area per lipid of all system, Radial Distribution Function of ARG139, ASP289 and all lipid head groups; Get Contact analysis of protein residue,

## Conflicts of interest

There are no conflicts to declare.

## Data availability

The data supporting this article have been included as part of the ESI.

## Author Contribution

SP: running simulations, data curation, formal analysis, investigation, methodology, software, writing—original draft preparation. PS: data curation, formal analysis, software. SS: system set-up, data curation, software, formal analysis. AD: conceptualization, project administration, resources, supervision, validation, and editing.

## Acknowledgements

The authors would like to thank the high performance computational facility provided by the Indian Institute of Technology, Jodhpur. S.P is thankful to Harikrishna Nair for the initial steps of system setup. A.D is thankful to the Department of Biotechnology Bio-RIDE scheme (Grant ID: DBT/PR65037/MRNAR/169/43/2026), Ministry of Science and Technology, Government of India for financial support and Param Rudra for computing facilities.

## Supporting Information

The supporting information is available: Area per lipid of all systems; Radial Distribution Function of ARG139, ASP289 and all lipid head groups; Get Contact analysis of protein residues; Time evolution of the C_*α*_-C_*α*_ distances Tyr101–Asp289 and Tyr101–Leu31 for all replicas at different 5-HT concentrations; Mutual information between the two distances, with block-permutation significance tests for individual replicas and pooled data. Root Mean Square Fluctuation and Tilt angle of protein helices at different 5-HT concentrations.

## Abbreviations

5-hydroxytryptamine: 5-HT: Serotonin
GPCR: G-Protein coupled receptor
Y4R: Neuropeptide Y4 receptor
PP: Pancreatic Polypeptide
BPM: Brain plasma membrane
POPC: 1-palmitoyl-2-oleoyl-sn-glycero-3-phosphocholine
SAPC: 1-Stearoyl-2-Arachidonoyl-sn-glycero-3-Phosphatidylcholine
SM: Sphingomyelin
PE: Phosphatidylethanolamine
PS: Phosphatidylserine
PIP2: Phosphatidylinositol 4,5-bisphosphate
CER: Ceramide
CHOL: Cholesterol

---

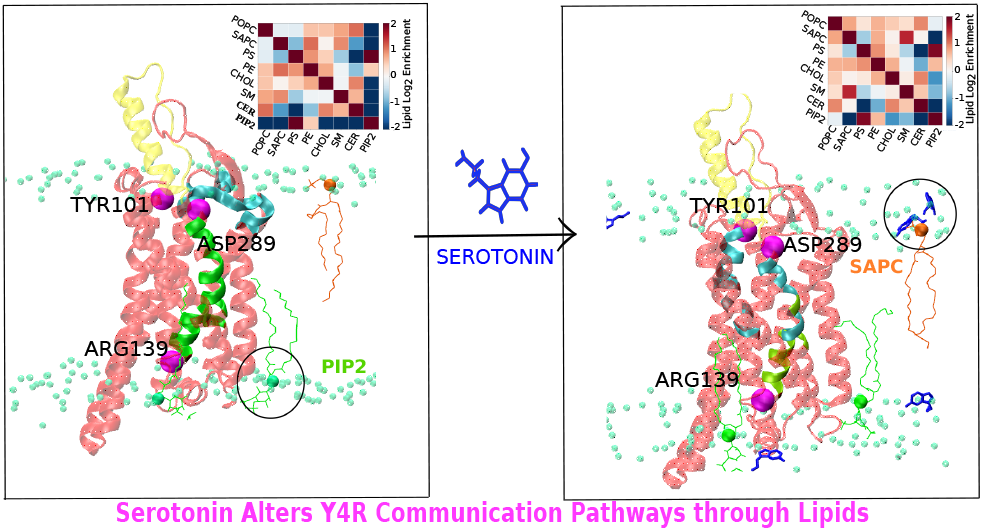

## Notes

### Competing Interest Statement

The authors have declared no competing interest.

