## Supplementary material for "Beyond Direct Binding: Serotonin Rewires Model Brain Plasma Membrane-Mediated Communication Pathways in Y4R": Area per lipid of all system, Radial Distribution Function of ARG139, ASP289 and all lipid head groups; Get Contact analysis of protein residue,

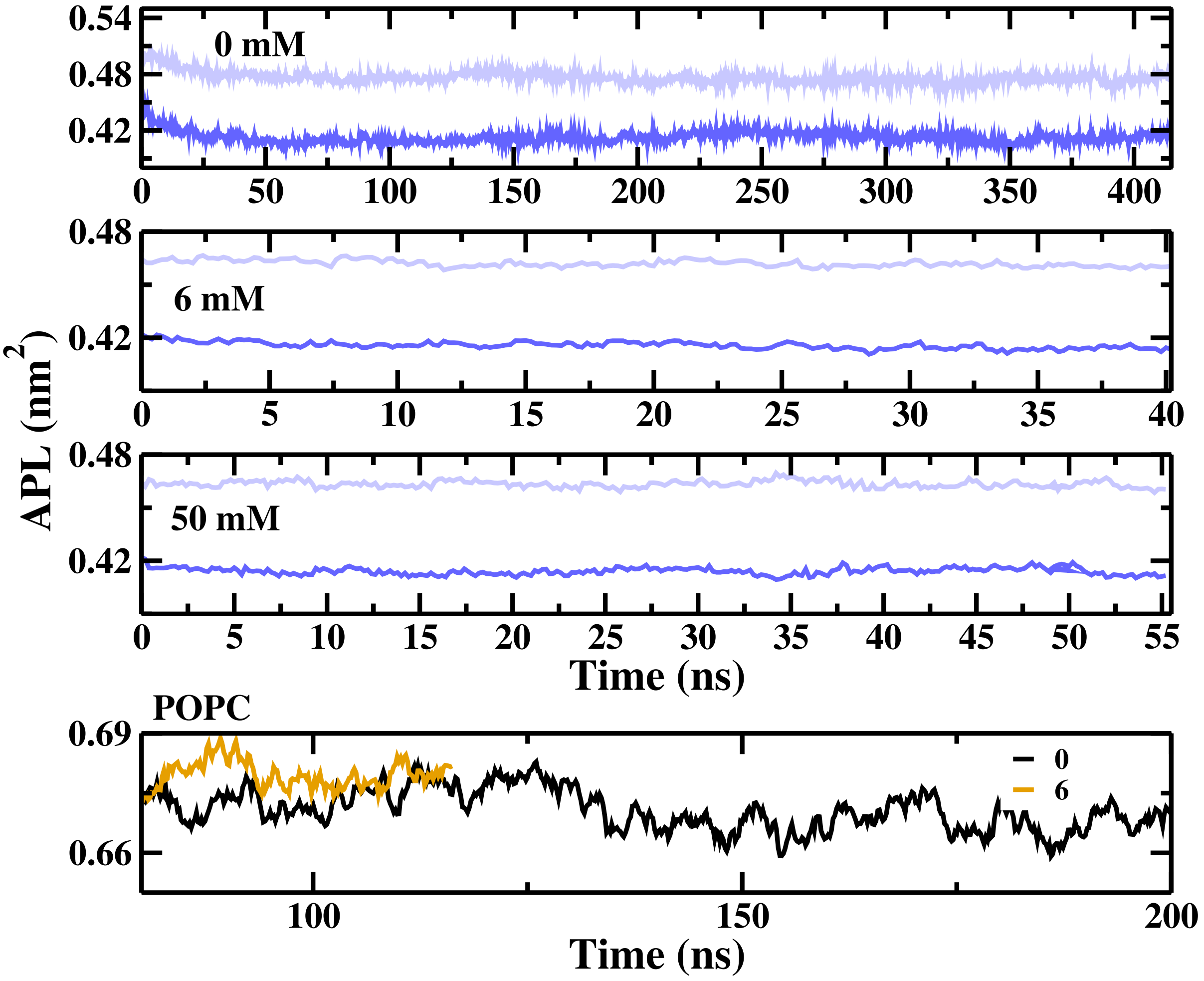


Figure S1: Convergence of the area per lipid during NPT equilibration runs, showing that all systems are well equilibrated.

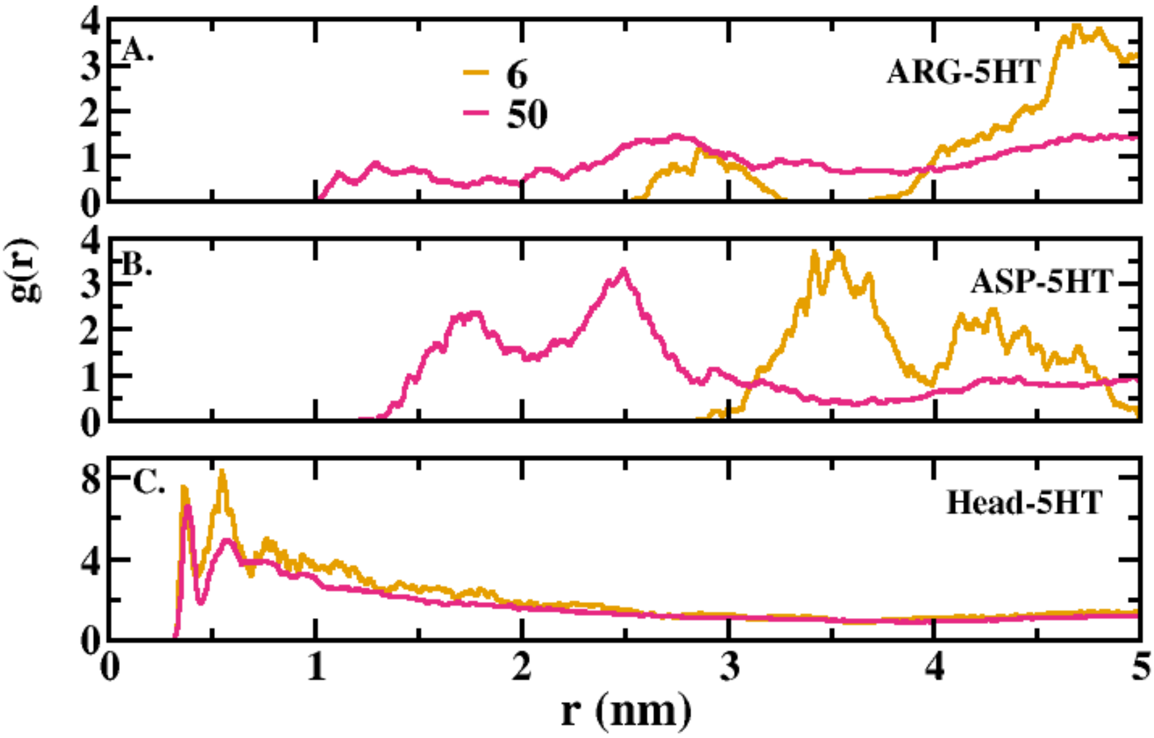


Figure S2: Radial distribution function of (A) ARG139, (B) ASP289, and (C) all lipid head groups with respect to 5-HT at all concentrations. This shows that 5-HT prefers to reside near the lipid head groups rather than the protein residues, experimentally identified to be important for PP binding (ASP289) and Y4R activation through G-protein coupling (ARG139).

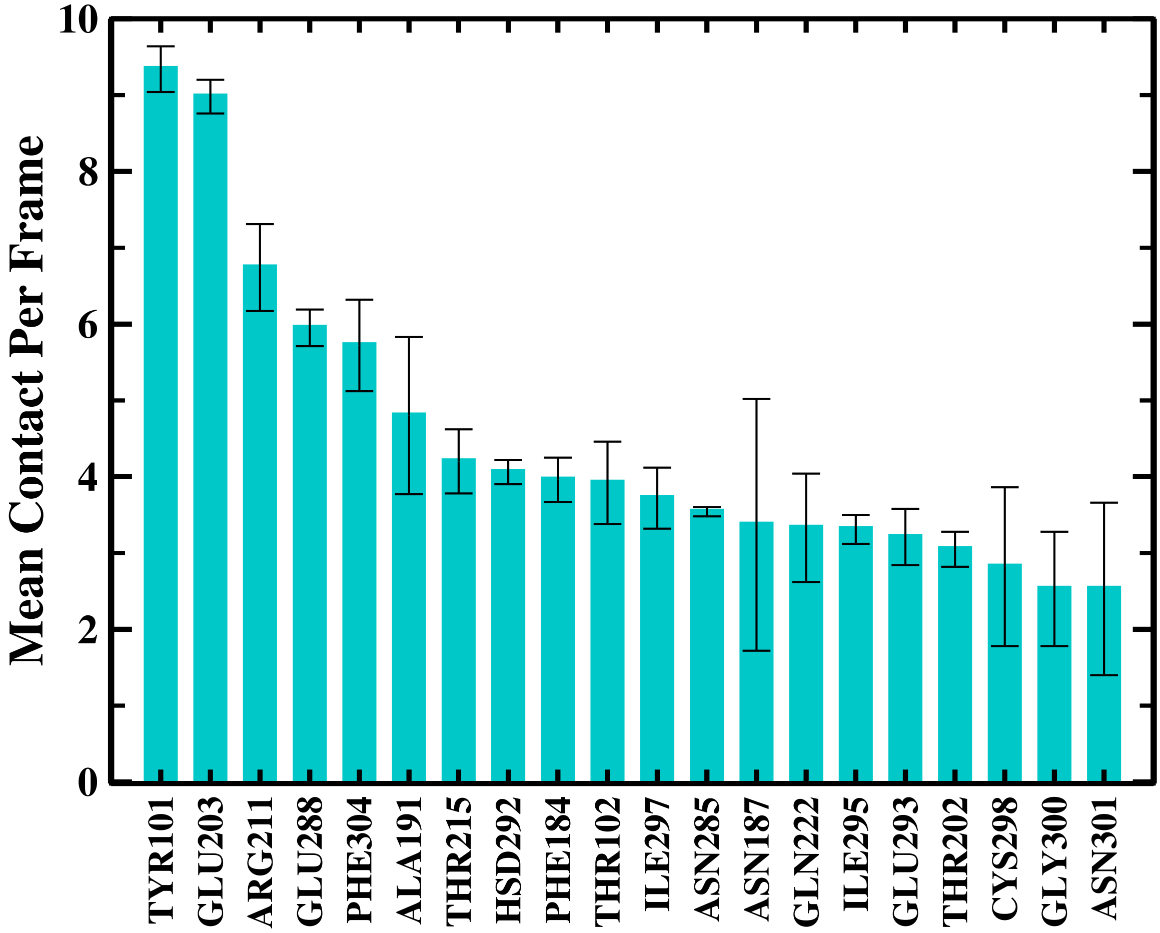


Figure S3: Average Contact of protein residues with ligand, which shows that TYR101 has the highest contact per frame to PP at 0 mM 5-HT concentration.

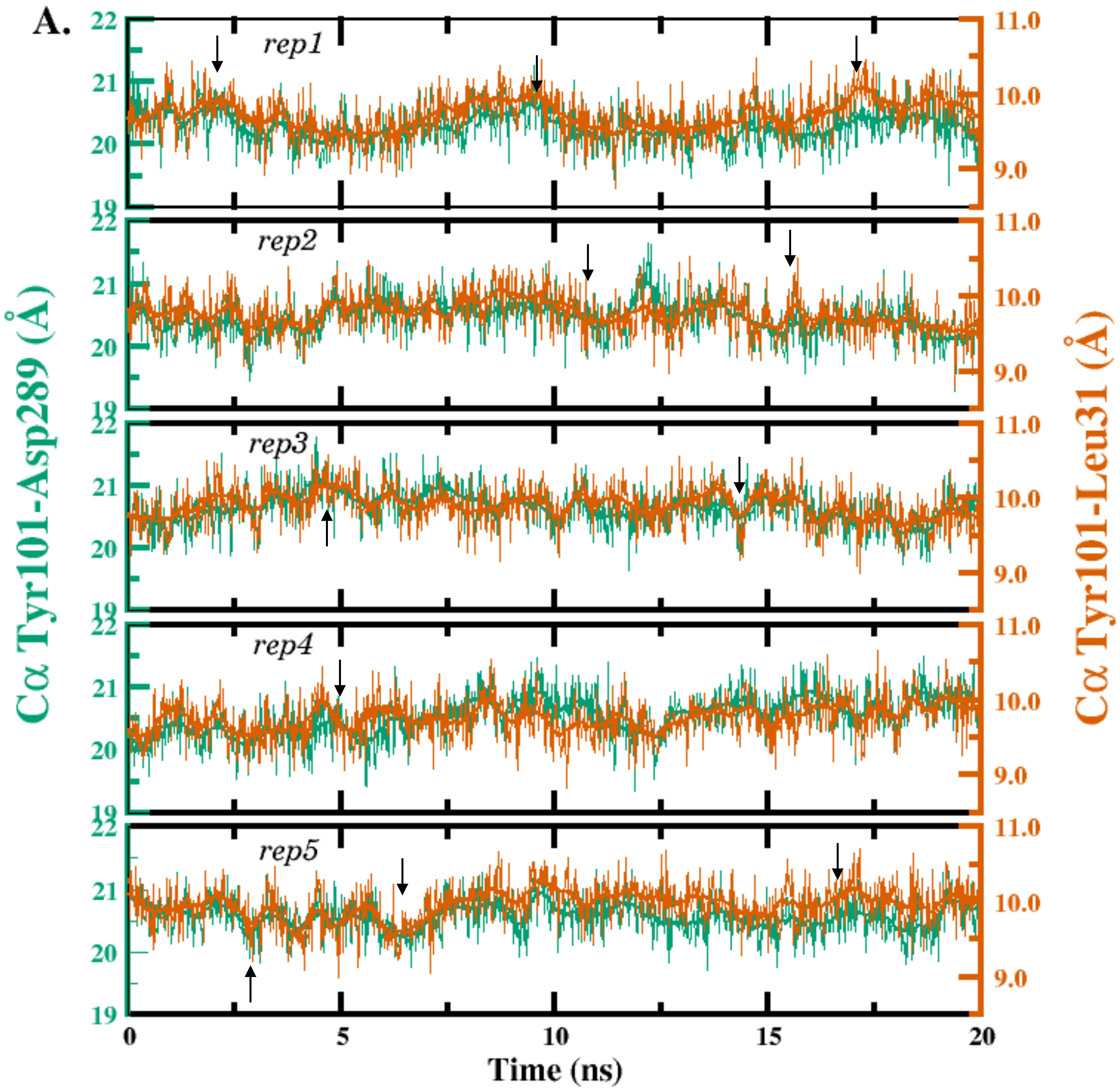


Figure S4: Time evolution of the $C_\alpha$-$C_\alpha$ distances Tyr101-Asp289 of Y4R (green, left axis) and Tyr101-Leu31 of PP (orange, right axis) over 20 ns for the five independent replicas (rep1-rep5) in the absence of 5-HT (0 mM). The black arrows indicate the correlation between Tyr101-Asp289 and Tyr101-Leu31.

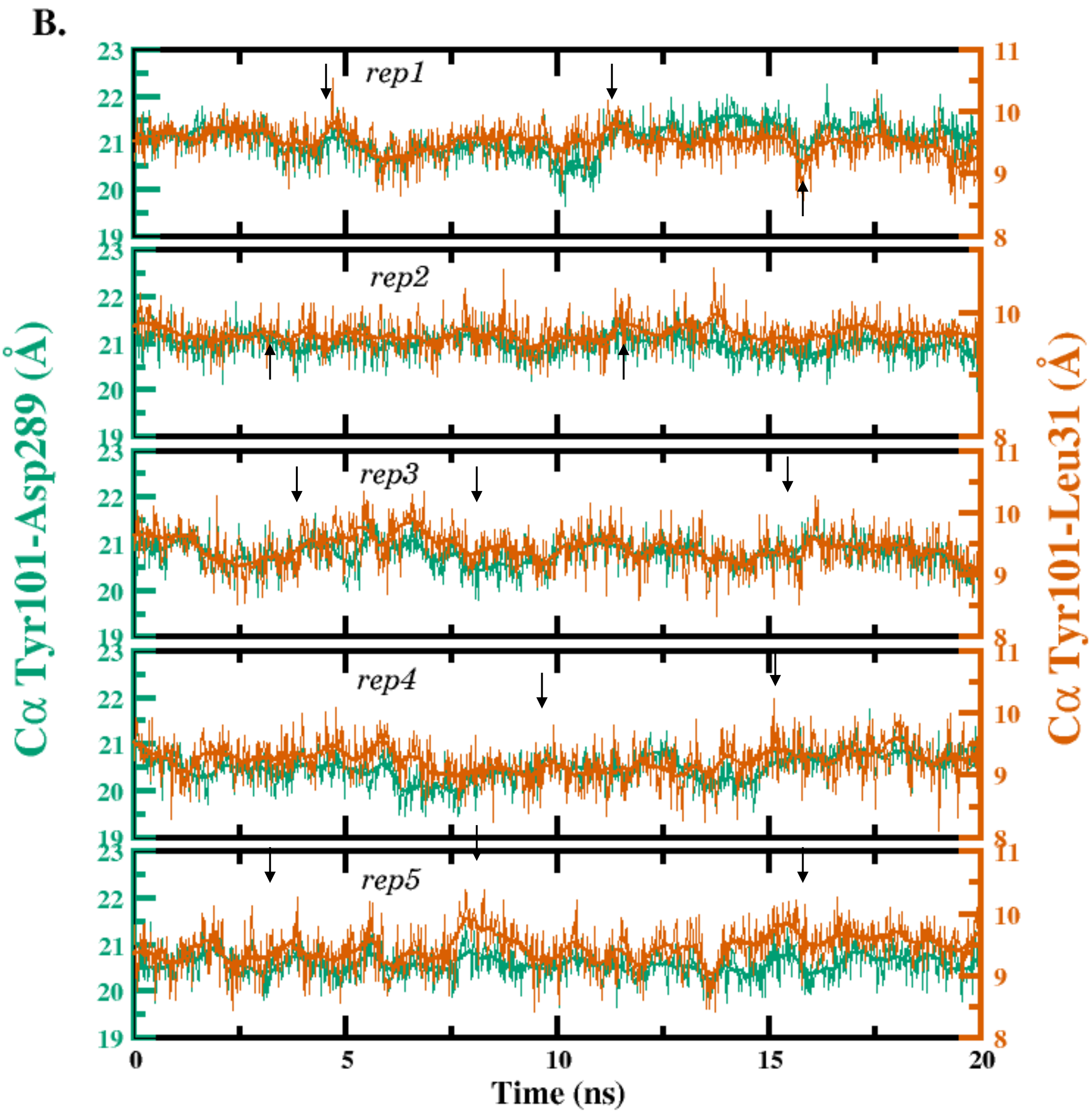


Figure S5: Time evolution of the $C_\alpha$-$C_\alpha$ distances Tyr101-Asp289 of Y4R (green, left axis) and Tyr101–Leu31 of PP (orange, right axis) over 20 ns for the five independent replicas (rep1-rep5) in the presence of 6 mM 5-HT. The black arrows indicate the correlation between Tyr101-Asp289 and Tyr101-Leu31.

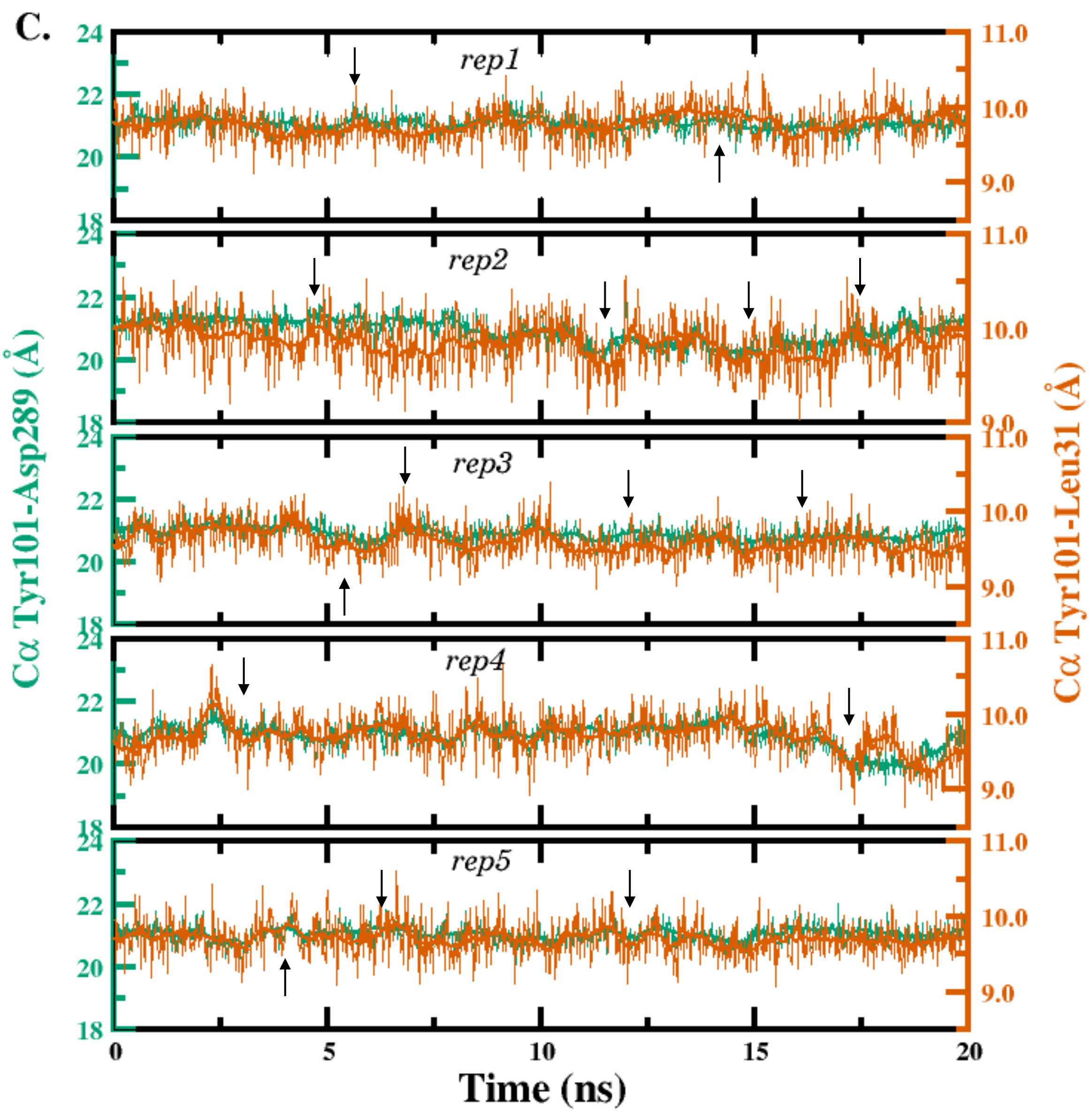


Figure S6: Time evolution of the $C_\alpha$-$C_\alpha$ distances Tyr101-Asp289 of Y4R (green, left axis) and Tyr101-Leu31 of PP (orange, right axis) over 20 ns for the five independent replicas (rep1–rep5) in the presence of 50 mM 5-HT. The black arrows indicate the correlation between Tyr101-Asp289 and Tyr101-Leu31.

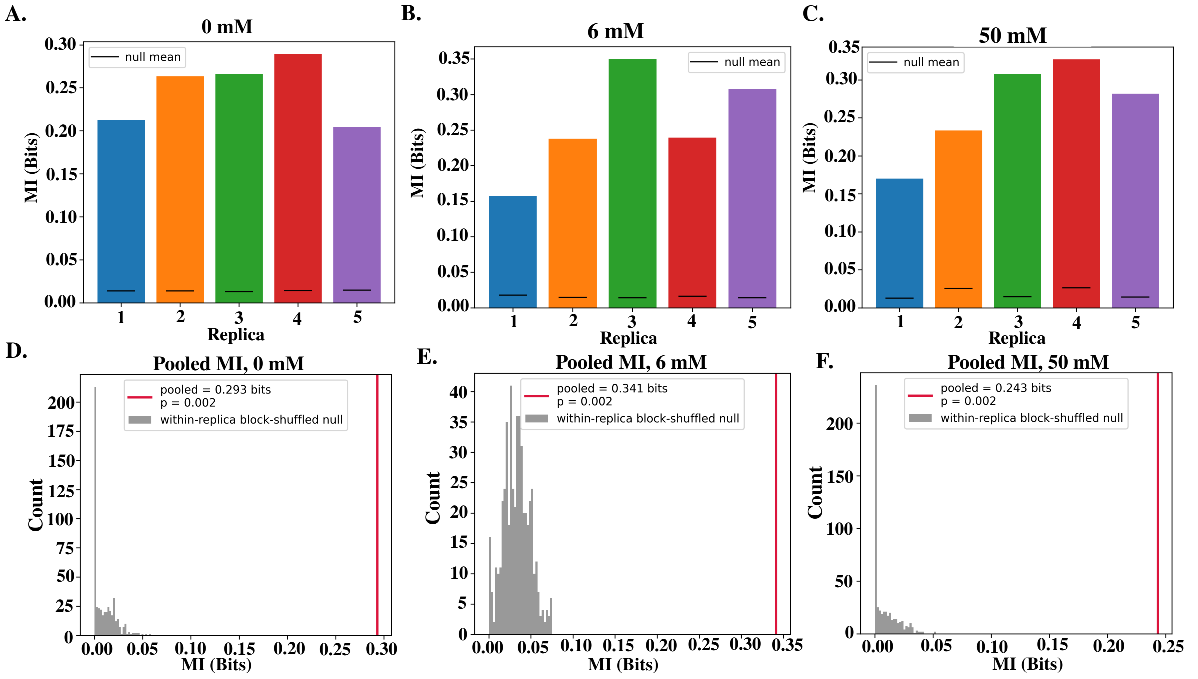


Figure S7: Mutual information (MI) between the Tyr101-Asp289 $C_\alpha$-$C_\alpha$ distance, $x(t)$, and the Tyr101–Leu31 $C_\alpha$–$C_\alpha$ distance, $y(t)$, at 0, 6 and 50 mM. **(A–C)** MI of each of the five independent replicas. The black horizontal line on each bar marks the mean MI of the corresponding block-permutation null distribution (500 permutations). In every replica the observed MI is far above the null mean. (D–F) Pooled MI over the five replicas at 0 mM (D), 6 mM (E) and 50 mM (F). The red vertical line shows the observed pooled MI (0.293, 0.341 and 0.243 bits, respectively), and the grey histogram shows the pooled null distribution obtained by shuffling $x(t)$ blocks within each replica while keeping $y(t)$ fixed. The observed pooled MI lies outside the null distribution in all three conditions ($p = 0.002$, the smallest value attainable with 500 permutations). MI is given in bits.

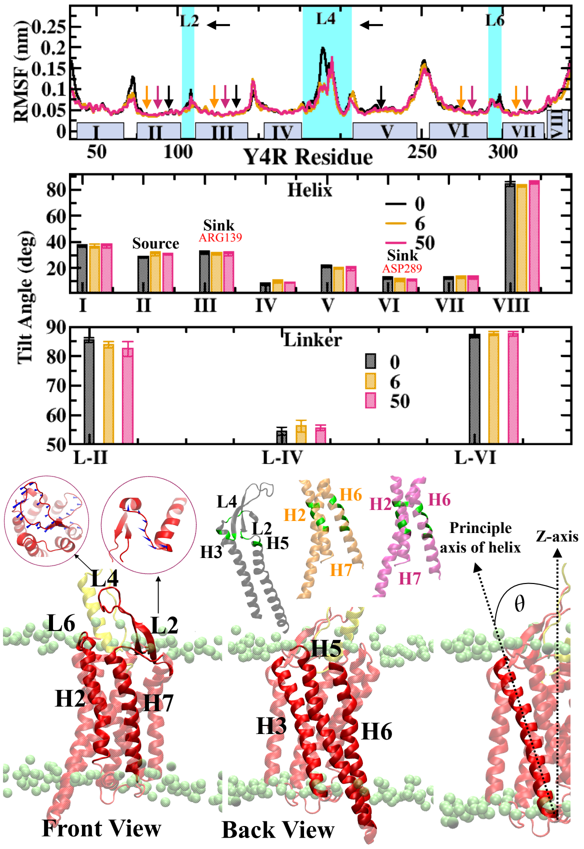


Figure S8: (A) Per-residue RMSF of Y4R, plotted against residue number for the transmembrane helices I–VII and the cytosolic helix VIII are indicated for the three conditions (black, 0; orange, 6; magenta, 50).In the same plot, the RMSF of extracellular loops L2, L4, and L6 are highlighted in cyan showing their elevated mobility. Colored arrows mark residues known to participate in receptor signaling, colored to match the condition (black, 0; orange, 6; magenta, 50) in which their involvement is assessed; each arrow is positioned indicating the helix and the linker to which that signaling residue belongs. (B) Average tilt angle (degrees, relative to the membrane normal/z-axis) of each helix (top) and the extracellular linker segments L-II, L-IV, and L-VI (bottom) are shown for the three conditions (black, 0; orange, 6; magenta, 50). The Source TYR101 belongs to transmembrane helix II and the Sink ARG139 and ASP289 belongs to helix III and VI are shown just to show the residues' tilt angle with respect to the addition of 5-HT. (C)Representative snapshots of Y4R embedded in the BPM bilayer (lipid headgroups shown as pale green spheres). Front and back views show the transmembrane helix bundle (H2, H3, H5, H6, H7) and loops L2, L4, and L6 which are responsible for the signal transduction. The two circular insets show porcupine plots of loops L2 and L4, in which blue sticks radiating the direction and relative magnitude of local backbone displacement. In the adjacent structural overlays, helices are colored black, orange, and magenta to represent conditions 0, 6, and 50, respectively, illustrating the conformational change of the H2–H3–H5–H6–H7 bundle across conditions; residues highlighted in green trace the signaling pathway and how its conformation shifts between conditions 0 and 6. The rightmost panel defines the helix tilt angle $\theta$ used in (B), measured between the principal axis of a transmembrane helix (dashed arrow) and the bilayer normal, or z-axis (dashed vertical line).
